# Mitotic phosphorylation of Lamin B1 rod domain by ULK1 and Aurora A/PLK1 promotes spindle function

**DOI:** 10.1101/2025.04.25.650690

**Authors:** María José Mendiburo, Claudia Kalus, Anna B. Hamacher, Arezoo Shahba, Niklas Berleth, Lena Berning, Yadong Sun, Padmashri Naren, Zehan Hu, Gary Kasof, Jörn Dengjel, Björn Stork

## Abstract

The coil-coil rod domain that mediates lateral assembly of lamin filaments has been shown by proteomic approaches to undergo phosphorylation, though the function of these modifications remains unknown. Here, we identify serine 210 (S210) within the Lamin B1 rod domain as a mitotic phospho-acceptor residue, regulated by the combined action of the autophagy-activating kinase ULK1 and the mitotic kinases Aurora A and PLK1. Using a phospho-specific antibody, we demonstrate that Lamin B1 phospho-S210 is enriched at the mitotic spindle and interacts with a network of proteins involved in spindle assembly and spindle pole focusing. Preventing S210 phosphorylation increases the number of cells with multipolar or shorter spindles and prolongs mitotic duration. Our findings indicate that mitotic phosphorylation of Lamin B1 at S210 within the rod domain is important for proper spindle organization and focusing during mitosis.

## Introduction

Lamins are type V intermediate filament proteins forming a protein meshwork located just beneath the inner nuclear membrane, known as the nuclear lamina. This structure was initially recognized as essential for the maintenance of the mechanical properties of the nucleus, but more recently has been found to also regulate genomic organization and gene expression (*1*).

Lamins share a similar structure, with a short N-terminal head, a central α-helical rod domain containing three coiled coil regions, and a C-terminal tail. Dimerization through parallel coiled-coil interactions of their rod domains and head-to-tail polymerization leads to the assembly of proto-filaments, which further assemble in an antiparallel manner to form filament structures. Unlike other lamin isoforms, Lamin B1 and Lamin B2, collectively called B-type lamins, are permanently anchored to the inner nuclear membrane. This targeting is facilitated by the post-translational farnesylation of a C-terminal CaaX motif, which probably promotes hydrophobic interactions at the inner nuclear membrane (*2*).

Beyond their structural role, B-type lamins regulate a wide range of nuclear processes, such as DNA replication, gene expression, cell cycle progression, and DNA repair (*3–10*). Maintaining proper levels of B-type lamins is crucial for cell physiology, as their dysregulation leads to pathologies affecting the central nervous system (*11–14*), poor prognosis in certain cancers (*15–19*), and permanent cell cycle arrest known as senescence (*3, 5, 20*). Notably, Lamin B1 degradation by autophagy, or “nucleophagy”, has been linked to the onset of senescence in response to oncogene expression and other genotoxic stressors (*6*).

During mitosis, the nuclear lamina disassembles in a process catalyzed by the phosphorylation of lamin head and tail domains by the kinases CDK1 and PKC (*21, 22*). Remarkably, after disassembly, membrane-attached B-type lamins do not simply vesicularize and evenly distribute throughout the cellular space, but rather organize into a membranous matrix surrounding the mitotic spindle (*23*). This discovery, made almost two decades ago in *Xenopus* egg extracts, was extended by the finding that this lamin B network both permeates and surrounds spindles in a RanGTP-dependent manner. Importantly, assembly of this matrix is necessary for proper mitotic spindle function, as illustrated by experiments in which the depletion of *Xenopus* LB3 (the only B-type lamin) or Lamin B1 and B2 in HeLa cells induced defects in the recruitment of spindle assembly factors to the matrix, resulting in poor spindle morphology. Computational models have proposed that this structure regulates microtubule polymerization and physically restricts microtubule movement, thereby improving spindle focusing (*24*). Although the exact mechanism for B-type lamin assembly into the spindle matrix has not been determined, the activity of microtubule end-directed motor proteins such as dynein and the kinesin Eg5 seems to be required (*25, 26*). A possible role of post-translational modifications on B-type lamins in this process has not yet been investigated.

The autophagy-activating kinase ULK1 or its close homolog ULK2 are components of the ULK1/2 complex, which is responsible for autophagy initiation in response to cellular stress. Notably, ULK1 also participates in a broad range of autophagy-independent processes (*27–32*). Particularly interesting is the existence of a nuclear pool of ULK1 in mouse embryonic fibroblasts (*28*), melanoma cells (*33*), and other human cell lines (*34*). Although the function of this nuclear ULK1 has thus far remained relatively unexplored, some studies have shown its involvement in PARP1 activation in response to oxidative stress (*28*) and binding to IRF1 to control the expression of IFNϒ-induced immunosuppressive genes (*33*).

Motivated by limited knowledge of nuclear ULK1 function and by Lamin B1’s status as an autophagy substrate under genotoxic stress, we investigated their potential link. We discovered that ULK1, with Aurora A and PLK1, phosphorylates Lamin B1 at serine 210 in the rod domain during mitosis. This phosphorylation marks Lamin B1 at the microtubule spindle, and its loss increases spindle focusing defects and slightly prolongs mitosis.

## Results

### ULK1 phosphorylates the Lamin B1 rod domain

To analyze whether ULK1 activity is linked to the regulation of Lamin B1 autophagy within the nucleus, we first confirmed the presence of ULK1 in this compartment by fractionating HeLa cells. Although considerably weaker than its cytoplasmic signal, ULK1 was detected as a clear band in nuclear extracts under homeostatic conditions (Fig. 1a). Next, due to the high insolubility of lamins, we tested whether ULK1 and Lamin B1 are in close proximity by an *in situ* proximity ligation assay (PLA) with fixed HeLa cells. This assay detects the proximity of two endogenous proteins in their natural context, producing a fluorescent signal when antibodies against them are within 40 nm of each other (*35*). We detected significantly more signals in the samples in which both anti-Lamin B1 and anti-ULK1 antibodies were present than in the samples in which only one of the antibodies was added as a negative control (Fig. 1b). Signals were localized mainly at the nuclear periphery, where Lamin B1 is highly enriched, suggesting that a pool of nuclear ULK1 is tethered to the lamina. Given that ULK1 is a serine-threonine kinase with a broad spectrum of reported substrates (*29, 31, 32, 36, 37*), we tested whether ULK1 can directly phosphorylate Lamin B1 by *in vitro* kinase assays. We incubated full-length purified Lamin B1 with ATP and either the kinase domain of ULK1 (amino acids 1-649) or with ULK3, another member of the ULK family. We included ULK3 in our initial analysis because its activity has been linked to the induction of cellular senescence (*38*), a state of stable cell cycle arrest that is strongly correlated with Lamin B1 degradation by nucleophagy (*6*). The reactions were analyzed by mass spectrometry to map potential phospho-acceptor residues in Lamin B1. After normalization to a Lamin B1-only control to remove phosphorylation sites present in the purified input protein, we detected two ULK1-dependent, three ULK3-dependent, and six ULK1- and ULK3-dependent phospho-sites (Table S1). From these, we considered relevant only class I sites with a localization probability higher than 0.75 (*39*). Notably, most of these sites were concentrated within the coiled-coil rod domain of Lamin B1 (Fig. 1c). To validate these results, we performed radioactive *in vitro* kinase assays with ULK1 and GST-fused Lamin B1 fragments. For this, GST-Lamin B1 head (amino-acids 1-34), a region comprising coil 1A, linker 1 and coil 1B (amino acids 35-215; from now on referred to as coil 1), a fragment comprising linker 2 and coil 2 (amino acids 216-386; from now on referred to as coil 2) or tail (amino acids 387-586) were incubated with GST-ULK1 (amino acids 1-649) in the presence of [ϒ-^32^P] ATP. We focused on ULK1-dependent phosphorylation of Lamin B1 due to its confirmed nuclear localization and the observed proximity between the two proteins. Indeed, coil 1 and coil 2 were readily phosphorylated by ULK1, consistent with the mass spectrometry results (Fig. 1d, left panel). To further identify the modified residues, we generated serine-to-alanine variants of each potential target residue. We repeated the kinase assay with these variants and found that serine 210 (S210) in coil 1 was the primary phospho-acceptor residue since this phosphorylation was completely absent in the phospho-abrogating S210A variant (Fig. 1d, right panel). For coil 2, the individual replacements of threonine 285 (T285) and S284 (originally detected as an ULK3-dependent site but included due to its proximity to T285) with alanine decreased the phosphorylation efficiency compared to that of wild-type (WT) coil 2 but did not abrogate the signal (Fig. S1). Combined replacement of S284, T285 and S288 with alanine further reduced coil 2 phosphorylation, suggesting that this cluster can be simultaneously targeted by ULK1 (Fig. S1).

**Figure 1:**
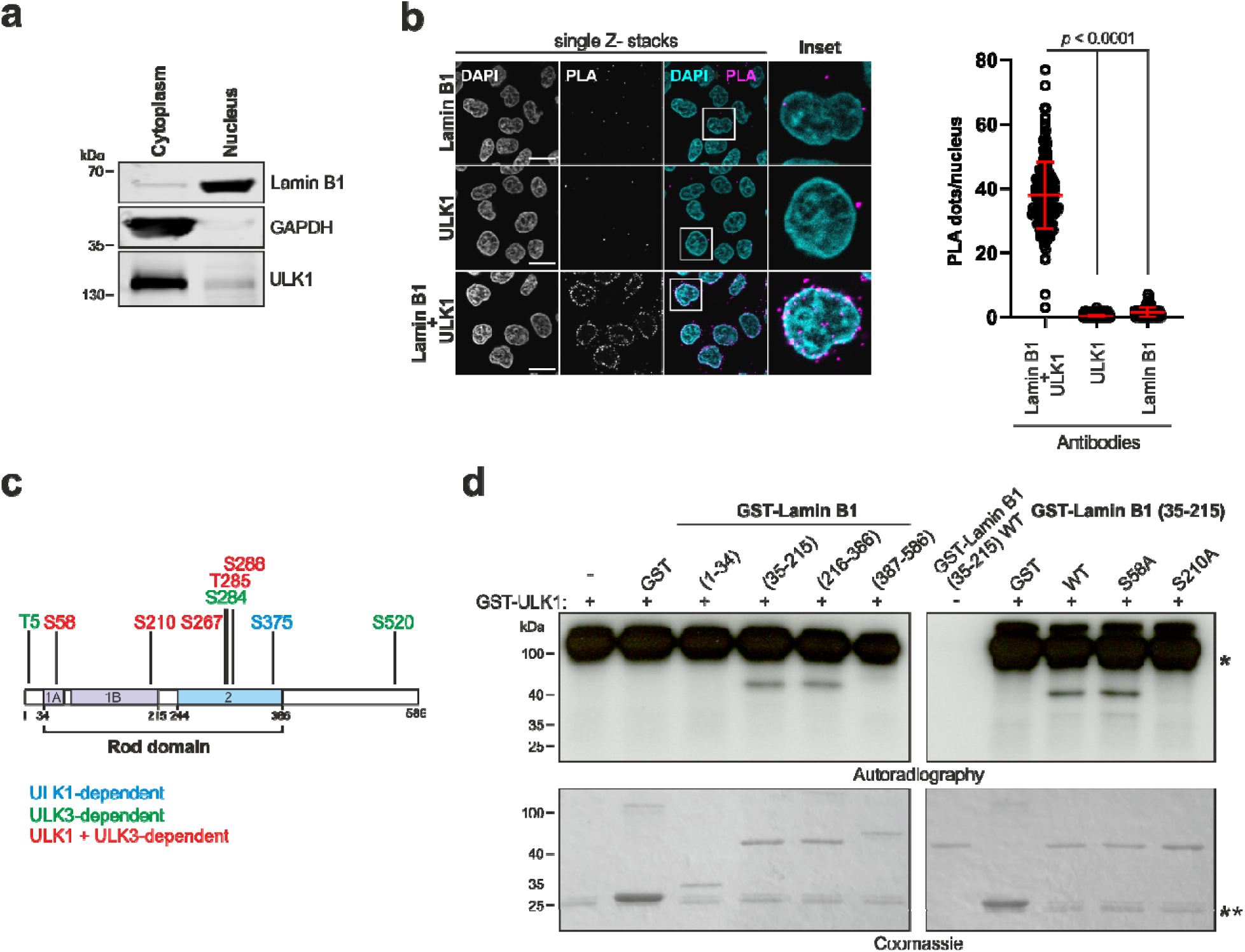
ULK1 phosphorylates Lamin B1 within the rod domain. **a)** HeLa cells were fractionated into cytoplasmic and nuclear fractions and immunoblotted with the indicated antibodies. **b)** Representative images (single z-stacks) of proximity ligation assay (PLA) with fixed HeLa cells co-incubated with Lamin B1 and ULK1 antibodies or treated with each antibody separately as controls. Nuclei are stained with DAPI. Scale bars, 10 μm. The right panel shows the quantification of the number of PLA signals per nucleus from one representative dataset out of three independent experiments. n_Lamin B1/ULK1_= 220, n_ULK1_= 224, n_Lamin B1_= 191 cells. *P* values were determined by one-way ANOVA with Tukey’s post hoc test. **c)** Schematic representation of the Lamin B1 protein domain organization (based on UniProt P20700), where the ULK1- and ULK3-dependent phospho-acceptor residues identified by mass spectrometry are depicted. Only sites with a localization probability higher than 0.75 (class I sites) are shown. **d)** Radioactive *in vitro* kinase assays with GST-ULK1 (1–649) and GST-Lamin B1 head, coil1, coil2 and tail fragments (left panel) or GST-Lamin B1 coil 1 WT, S58A or S210A variants (right panel). After incubation with [ϒ-^32^P] ATP in the presence of kinase buffer, reactions were separated via SDS‒PAGE and exposed to film (autoradiography). * indicates GST-ULK1 auto-phosphorylation, ** indicates free GST.

We focused on Lamin B1 S210 because of ULK1’s clear dependency on its phosphorylation. S210 phosphorylation appears to occur naturally in cells, as it has been detected in several high-throughput mass spectrometry studies across both stem cells and various cancer cell lines (https://www.phosphosite.org/siteAction.action?id=18011426). Analysis of the sequence around S210 (EDLEFRKS_210_MYEEEIN) revealed good conformity to the published ULK1 consensus motif, with hydrophobic residues at positions -3, +1, and +2 relative to the phosphoserine (*37*).

### Lamin B1 phosphorylated at S210 localizes dynamically to the mitotic spindle

To analyze the physiological relevance of S210 phosphorylation, we generated a S210 phospho-specific antibody in collaboration with Cell Signaling Technology. We initially tested the antibody by co-transfecting ULK1 and Lamin B1-myc coding vectors into 293T cells and analyzing Lamin B1 phospho-S210 (pS210) levels by immunoblotting. While single transfections led to a modest increase in the Lamin B1 pS210 signal, a substantial accumulation was observed when both vectors were co-transfected (Fig. S2a), supporting the antibody’s specificity for phosphorylated Lamin B1 and confirming ULK1 as an upstream kinase for this modification.

Since most of the Lamin B1 phosphorylation events known to date are related to the disassembly of the nuclear lamina at mitotic entry, we analyzed whether Lamin B1 S210 phosphorylation is a mitotic event by synchronizing HeLa cells with RO-3306 (a CDK1 inhibitor that arrests cells in G2/M) and subsequently releasing them into fresh media for different time points. A signal at approximately 70 kDa (slightly greater than that of total Lamin B1, which runs at approximately 66-68 kDa) was detected; this signal steadily increased within the first 60 minutes after RO-3306 washout and subsequently decreased (Fig. 2a), which correlated with an increase in the mitotic marker H3 phospho-S10 (H3 pS10). Interestingly, mitotic arrest induced by drugs affecting microtubule stability had distinct effects on the extent of phosphorylation; while microtubule stabilization with paclitaxel induced an 11.5-fold higher phospho-signal compared to the level observed in non-treated cells, depolymerization with nocodazole led to only a 5.0-fold increase (Fig. 2b), suggesting the existence of segregated pools of Lamin B1 that depend differentially on microtubule integrity for efficient phosphorylation at S210. A similar effect was observed in HCT116 cells (Fig. S2b). As expected, western blot analysis of Lamin B1 KO cells synchronized by RO-3306 treatment and released for 60 minutes revealed no signal for neither Lamin B1 pS210 nor total Lamin B1 (Fig. S2c), and treatment with with λ-phosphatase after synchronization by RO-3306 or arrest with paclitaxel induced a complete abrogation of the signal, indicating that the antibody is phospho-specific (Fig. S2d). To confirm that the target is phosphorylated S210 we purified GFP-ULK1 (WT or kinase-dead (KD; D164A)) from Flp-In™ T-Rex™ 293 cells (*40*) and performed *in vitro* kinase assays with GST-Lamin B1 coil1 (35–215) WT or carrying a S210A mutation. After the kinase reaction, proteins were separated by SDS‒PAGE and immunoblotted with the anti-Lamin B1 pS210 antibody. Although a background band corresponding to GST-Lamin B1 coil1 (35–215) WT without kinase (incubated with GFP as a negative control) was visible in this *in vitro* context, the signal was markedly enhanced when treated with GFP-ULK1 WT and was below the background level in GST-Lamin B1 coil1 (35–215) S210A incubated with GFP-ULK1 WT (Fig. S2e), confirming the greater affinity of the antibody for phosphorylated S210.

**Figure 2:**
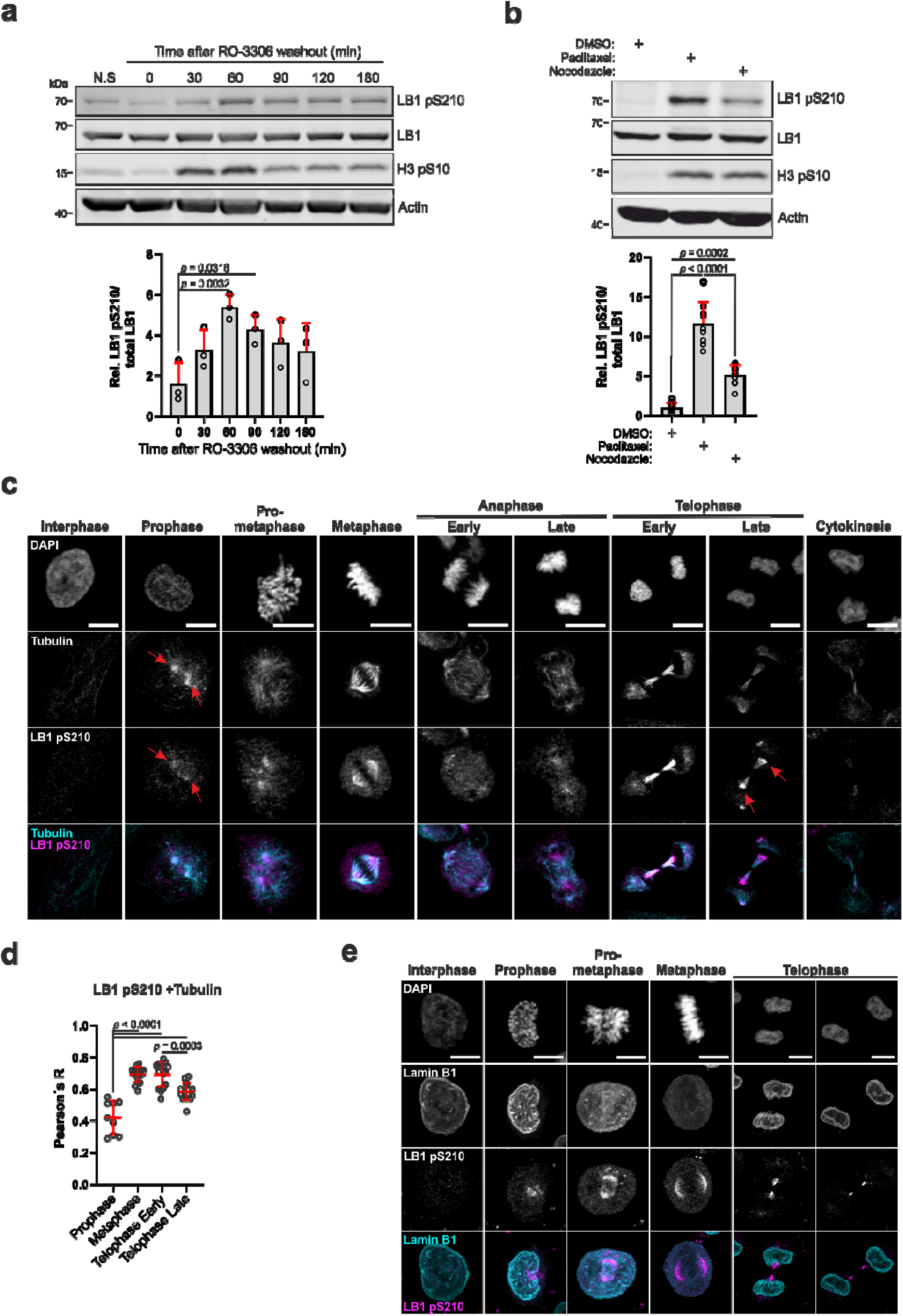
Lamin B1 phosphorylated at S210 localizes dynamically to the mitotic spindle. **a)** Representative immunoblots of HeLa cells arrested in G2/M phase with RO-3306 and released into fresh medium for the indicated times. N.S. = not synchronized. LB1 = Lamin B1. Lower panel: Quantification of the densitometry of Lamin B1 phospho-S210 bands standardized to the corresponding total Lamin B1 signal and normalized to time point 0. The data are presented as the means + SDs from three independent experiments. *P* values were determined by one-way ANOVA with Dunnett’s post hoc test. **b)** Representative immunoblots of HeLa cells arrested with 100 nM paclitaxel or 100 ng/ml nocodazole for 18 hr. LB1 = Lamin B1. Lower panel: Quantification of the densitometry of Lamin B1 pS210 bands standardized to the corresponding total Lamin B1 signal and normalized to the DMSO-treated sample. The data are presented as the means + SDs from nine independent experiments. *P* values were determined by one-way ANOVA with Tukey’s post hoc test. The actin immunoblots corresponding to the H3 pS10 signal in a) and b) are shown in the Source Data file. **c)** Representative images (z-stack projections) of HeLa cells synchronized in G1 by double-thymidine block and processed for immunofluorescence with anti-Lamin B1 pS210 and anti-α-Tubulin antibodies. DNA was stained with DAPI. The red arrows in the prophase panel indicate the position of centrosomes. In the late telophase image, the arrows show the segregation of the Lamin B1 phospho-S210 signal toward the nuclei and away from the α-Tubulin-stained midbody. Scale bars, 10 μm. **d)** Quantification of the correlation (Pearson’s R coefficient) between the Lamin B1 pS210 and the α-Tubulin signals in c). The data shown are pooled from two independent experiments, lines represent the means ± SDs. n_Prophase_ = 9, n_Metaphase_ = 16, n_Telophase early_ = 15, n_Telophase late_ = 16 cells. *P* values were determined by one-way ANOVA with Tukey’s post hoc test. **e)** Representative images (z-stack projections) of HeLa cells synchronized by double-thymidine block and processed for immunofluorescence with anti-Lamin B1 and anti-Lamin B1 pS210 antibodies. DNA was stained with DAPI. Scale bars, 10 μm.

Next, we examined the localization of Lamin B1 pS210 by co-staining with an anti-Tubulin antibody by immunofluorescence. In line with pS210 being a mitotic phosphorylation, we did not detect a signal in cells in interphase, indicating that the phospho-antibody does not cross-react with unphosphorylated Lamin B1 at the nuclear lamina. Intriguingly, when analyzing cells in mitosis, we unexpectedly observed that Lamin B1 pS210 was concentrated in regions near microtubules and the mitotic spindle. The signal emerged early in prophase close to centrosomal regions, as visualized by a focused Tubulin signal (Fig. 2c, red arrows in the prophase panels). In later mitotic stages, Lamin B1 pS210 localized mainly at the spindle poles in metaphase and translocated toward the central spindle at anaphase, where it accumulated during telophase. In late telophase, it segregated away from Tubulin toward the re-forming nuclei (Fig. 2c, red arrows in the late telophase panel) and finally disappeared before cytokinesis was complete. Measurement of the Pearson’s R coefficient in individual images from these cells revealed a progressive increase in the correlation between the two signals in metaphase and early telophase, which decreased during late telophase (Fig. 2d). This dynamic localization was also detected in mitotic HCT116 cells (Fig. S2f), where we additionally confirmed the phospho-specificity of the antibody in immunofluorescence by treatment with λ-phosphatase prior to incubation with anti-Lamin B1 pS210 (Fig. S2g). Of note, we detected only a partial overlap between total Lamin B1 and Lamin B1 pS210 signals by immunofluorescence throughout mitosis (Fig. 2e). pS210 is detectable in prophase close to centrosomes while total Lamin B1 is still localized at the nuclear envelope, and remains associated with microtubules at the midbody arms until after Lamin B1 is targeted to the re-forming nuclear envelope.

Taken together, our results point toward the existence of a Lamin B1 pool that becomes phosphorylated at S210 at mitotic entry and associates with the microtubule spindle throughout mitotic progression. Although the presence of B-type lamins at the mitotic spindle has been recognized for some years, to our knowledge, this is the only post-translational modification known so far to be specifically enriched at this region.

### ULK1 contributes to Lamin B1 S210 phosphorylation

Next, we investigated whether mitotic Lamin B1 phosphorylation at S210 depends on ULK1 activity in cells. Due to their high sequence similarity across the entire protein, particularly within their kinase domains, ULK1 and its closely related homolog ULK2 are considered to share substrate specificity for several autophagy proteins (*41*). To avoid possible functional compensation of ULK2 in the absence of ULK1, we generated ULK1 and ULK2 double-knockout (ULK1/2 DKO) HeLa cells by CRISPR-Cas9 editing.

Synchronization experiments with WT and ULK1/2 DKO cells revealed a mild but significant reduction in the abundance of the Lamin B1 pS210 signal according to western blot analysis in the absence of ULK1/2. WT cells displayed a 6.0- and 5.4-fold increase in the S210 phospho-signal relative to that in the G2/M-arrested cells (no release from RO-3306) 60 and 90 minutes after RO-3306 washout, respectively, whereas these values reached only 3.2- and 2.6-fold in the ULK1/2 DKO cells (Fig. 3a). We observed a similar effect upon paclitaxel treatment, in which WT cells exhibited a 12.7-fold increase in the Lamin B1 pS210 signal relative to that in non-treated cells, while in ULK1/2 DKO cells the increase was 8.0-fold (Fig. 3b). To analyze these effects via a complementary approach, we depleted ULK1 by siRNA in HeLa WT cells and repeated the paclitaxel treatment. The absence of ULK1 was sufficient to reduce the intensity of the Lamin B1 pS210 band to approximately half of the level observed in cells treated with non-targeting siRNA (Fig. 3c), supporting the results obtained with DKO cells. In line with these observations, immunofluorescence staining of WT and DKO cells with anti-Lamin B1 pS210 revealed a more diffuse localization and slightly lower signal intensity at the mitotic spindle of DKO metaphase cells (Fig. S3a). We additionally noted that ULK1 becomes enriched in the area where microtubules organize around the centrosome during early prophase (Fig. S3b, red arrow) and later distributes evenly throughout the cytoplasm. This localization also correlates with the region where Lamin B1 phosphorylation at S210 is first detected (Fig. 2c), suggesting that ULK1 may contribute to early Lamin B1 phosphorylation locally. Due to the lack of a reliable mouse anti-ULK1 antibody required for PLA with our rabbit anti-Lamin B1 pS210, we reconstituted ULK1/2 DKO cells with flag-myc-ULK1 or an empty vector as a control, and performed the assay using mouse anti-Flag and the phospho-antibody. We detected PLA signals in cells in prophase, pro-metaphase and metaphase which were absent in cells reconstituted with the empty vector (Fig. 3d). In line with this, analysis of endogenous ULK1 and total Lamin B1 in WT and ULK1/2 DKO cells (as negative control) revealed significantly more PLA signals in mitotic WT cells (Fig. S3c). Notably, in this case the signal was slightly higher in cells in prophase than in pro-metaphase and metaphase and concentrated in the nuclear area in contrast to the more cytoplasmic localization observed in the reconstituted setup (Fig. 3d). These results indicate that ULK1/Lamin B1 proximity generally increases in the nuclear area during prophase, whereas the phosphorylation of S210 likely occurs in a region adjacent to the nucleus after nuclear envelope breakdown.

**Figure 3:**
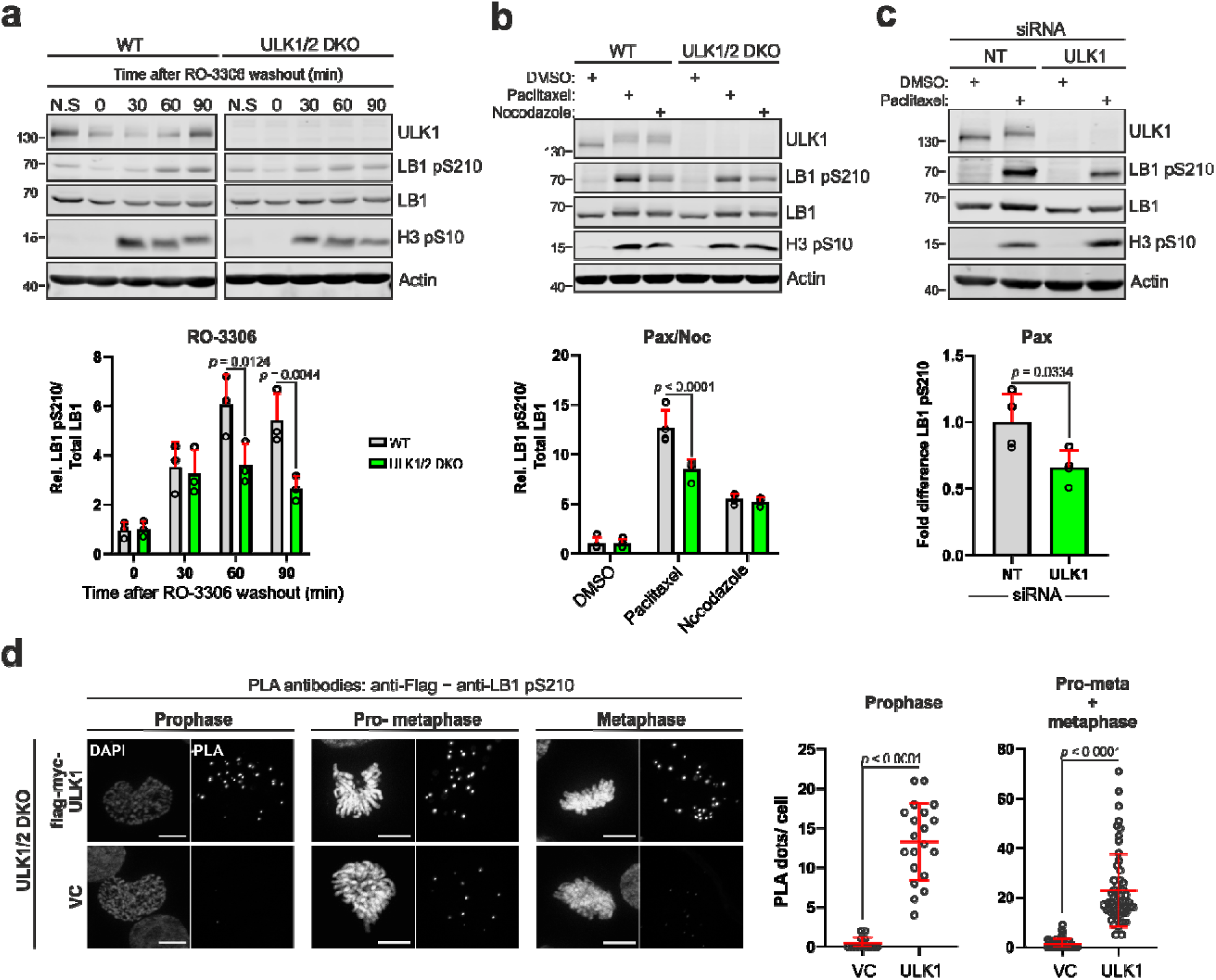
ULK1 partially contributes to Lamin B1 S210 phosphorylation. **a)** Representative immunoblots of WT and ULK1/2 DKO cells arrested in G2/M with RO-3306 and released into fresh medium for the indicated times. N.S = not synchronized. Lower panel: Quantification of Lamin B1 pS210 densitometry. The data are presented as the means + SDs from three independent experiments. *P* values were determined by two-way ANOVA with Sidak’s post hoc test. **b)** Representative immunoblots of WT and ULK1/2 DKO cells arrested with paclitaxel or nocodazole. The up-shift of ULK1 in mitosis is due to its hyper-phosphorylation by CDK1 (*76*). Lower panel: Quantification of Lamin B1 pS210 densitometry. The data are presented as the means + SDs from three independent experiments. *P* values were determined by two-way ANOVA with Sidak’s post hoc test. **c)** Representative immunoblots of WT cells treated with ULK1 or non-targeting (NT) siRNA and exposed to paclitaxel or DMSO. Lower panel: Quantification of the Lamin B1 pS210 band densitometry. The data are presented as the means + SDs from four independent experiments. The *P* value was determined by two-way ANOVA with Sidak’s post hoc test. The actin immunoblots corresponding to the H3 pS10 signal in a), b) and c) are shown in the Source Data file. **d)** Representative images of PLA assays with anti-Flag and anti-Lamin B1 pS210 antibodies in mitotic HeLa ULK1/2 DKO cells reconstituted with flag-myc-ULK1 or vector control (VC). Scale bars, 10 μm. The graphs on the right show the quantification of PLA signals per individual mitotic cell. Pooled data from two independent experiments are shown, lines represent the means ± SDs. n_VC_Prophase_ = 14, n_flag-myc-ULK1_Prophase_ = 20, n_VC_Pro-meta/metaphase_ = 57, n_flag-myc-ULK1_Pro-meta/metaphase_ = 56 cells. *P* values were determined by Mann-Whitney test.

Our data suggest that ULK1 locally regulates Lamin B1 phosphorylation at S210 at mitotic entry. However, given the residual phosphorylation in observed ULK1/2 depleted cells, it is likely that this activity is complemented by other kinases targeting this site.

### PLK1 and Aurora A also phosphorylate Lamin B1 S210 during mitosis

To identify other potential regulatory kinases, we turned our attention to Aurora kinases A (AurA) and B (AurB) and Polo-like kinase 1 (PLK1), which are known to cooperatively regulate a plethora of mitotic processes (*42*). Both AurA and PLK1 co-localize and are active at centrosomes during prophase, the stage at which we observed the emergence of the Lamin B1 pS210 signal by immunofluorescence. We co-stained Lamin B1 pS210 with either AurA or PLK1 in WT HeLa cells and observed the localization of both kinases at punctate structures in prophase cells, likely corresponding to centrosomes (Fig. S4a, red arrowheads). Interestingly, Lamin B1 pS210 was detected in the area surrounding the centrosome-localized kinases but did not overlap with them (Fig. S4a, insets). During metaphase, the Lamin B1 pS210 signal strongly correlated with AurA at the spindle poles, whereas there was a lower co-localization with PLK1 (Fig. S4a, graph). PLA analysis of cells in prophase, pro-metaphase and metaphase with antibodies directed against each kinase and Lamin B1 pS210 revealed a significant enrichment of signal compared with each single-antibody control for both kinases (Fig. 4a). A similar trend was observed when using antibodies against AurA or PLK1 and total Lamin B1 (Fig. S4b), although in this setting the signal detected for PLK1/Lamin B1 was significantly higher than for AurA/Lamin B1.

**Figure 4:**
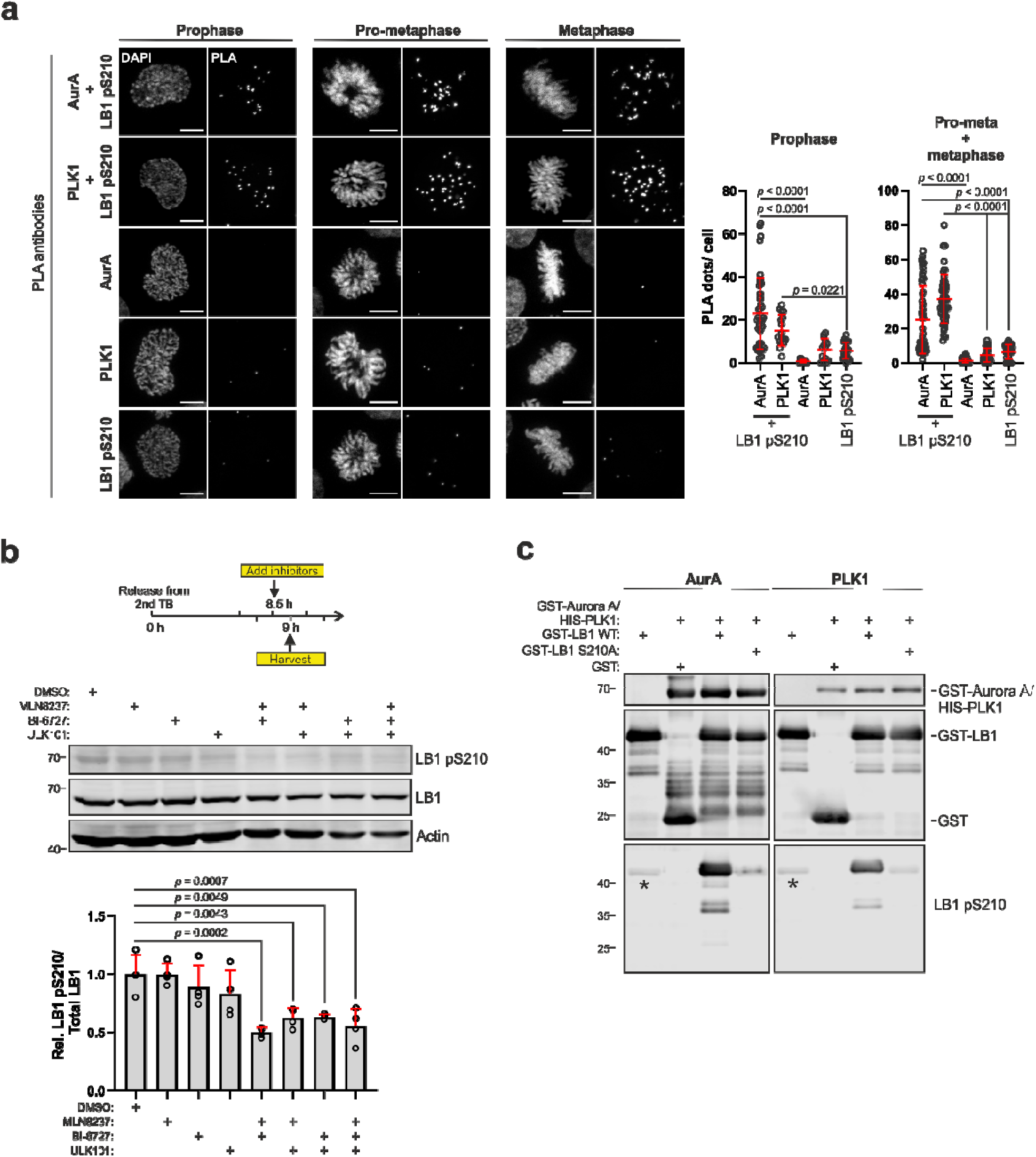
PLK1 and Aurora A cooperatively phosphorylate Lamin B1 S210 during mitosis. **a)** Representative images of PLA assays with the indicated antibodies in synchronized HeLa cells. Scale bars, 10 μm. The graphs show the quantification of PLA signals per individual mitotic cell. Pooled data from two to three independent experiments are shown, lines represent the means ± SDs. n_LB1 pS210/AurA_Prophase_= 44, n_LB1 pS210/PLK1_Prophase_= 17, n_AurA_Prophase_= 13, n_PLK1_Prophase_= 14, n_LB1 pS210_Prophase_= 24, n_LB1 pS210/AurA_Pro-meta/metaphase_= 64, n_LB1 pS210/PLK1_ Pro-meta/metaphase_= 57, n_AurA_ Pro-meta/metaphase_= 55, n_PLK1_ Pro-meta/metaphase_= 41, n_LB1 pS210_ Pro-meta/metaphase_= 41 cells. *P* values were determined by one-way ANOVA with Tukey’s post hoc test. **b)** Schematic representation of the experimental design for the treatment with different kinase inhibitors (upper panel). Middle panel shows immunoblot with the indicated antibodies. The graph below shows the quantification of the densitometry of Lamin B1 pS210 bands standardized to the corresponding total Lamin B1 signal and normalized to the DMSO-treated sample. The data are presented as the means + SDs from four independent experiments. *P* values were determined by one-way ANOVA with Dunnett’s post hoc test. **c)** Non-radioactive kinase assay with GST-Lamin B1 (35–215) WT or S210A and either GST-Aurora A (left panel) or HIS-PLK1 (right panel). After the kinase reaction, the samples were immunoblotted with the indicated antibodies. Asterisks in the anti-Lamin B1 pS210 blots indicate signals present in GST-Lamin B1 (35–215) WT without kinase.

Next, we tested whether pharmacological inhibition of each kinase, including ULK1, affects Lamin B1 pS210 levels. Given that AurA and PLK1 activities are crucial for mitotic entry (*43–45*), we synchronized the cells by double-thymidine block, added the inhibitors for AurA (MLN8237), PLK1 (BI-6727), or ULK1 (ULK101) 8.5 hr after the second release, and harvested them 30 min later (Fig. 4b, upper scheme). Western blot analysis revealed that treatment with each kinase inhibitor alone resulted in no significant difference in the level of Lamin B1 pS210 relative to that in DMSO-treated controls, whereas the combined blockade of AurA and PLK1 activities significantly decreased Lamin B1 S210 phosphorylation by half (Fig. 4b). Co-inhibition of AurA or PLK1 and ULK1 led to a reduction in the Lamin B1 pS210 level compared to each treatment, but the triple inhibition of AurA, PLK1, and ULK1 did not further enhance this outcome. These results suggest that AurA, PLK1, and ULK1 can redundantly phosphorylate Lamin B1 S210 under our experimental conditions.

PLK1 and AurA inhibition profoundly affect bipolar spindle assembly (*44, 46, 47*), which may, in turn, indirectly influence the abundance of Lamin B1 pS210. Thus, we tested whether PLK1 and AurA can directly phosphorylate Lamin B1 S210 by *in vitro* kinase assays and subsequent immunoblotting with the anti-Lamin B1 pS210 antibody. We detected a strong signal in the samples where each kinase was incubated with GST-Lamin B1 coil1 (35–215) WT but not the S210A variant, indicating that both kinases can indeed target this site (Fig. 4c). Altogether, these results support a relevant role for AurA and PLK1 in the mitotic phosphorylation of Lamin B1 at S210 in addition to ULK1.

### Lamin B2 S224 is targeted by AurA and PLK1 but not by ULK1

Lamin B1 and B2 are the only members of the mammalian B-type lamin family, sharing a 60% identical amino acid sequence. Alignment of the Lamin B1 and B2 regions surrounding S210 revealed conservation of S210 in Lamin B2 (S224; Fig. 5a), prompting us to test whether Lamin B2 is also a substrate for ULK1, AurA, and PLK1. To this end, we performed radioactive kinase assays with each kinase and GST-Lamin B2 coil1 (49–229) WT or S224A. The WT fragment was phosphorylated by both AurA and PLK1, whereas the inclusion of the S224A fragment or the addition of inhibitors for AurA (TC-S7010) and PLK1 (BI-6727) completely abolished the phosphorylation (Fig. 5b). To test whether ULK1 also phosphorylates Lamin B2 at S224, we performed kinase assays with GFP-ULK1, either WT or kinase-dead (KD), and GST-Lamin B2 coil1 (49–229) WT or S224A, including the GST-Lamin B1 coil 1 (35–215) fragment as a positive control for the reaction. As previously observed, incubating GFP-ULK1 WT with GST-Lamin B1 coil 1 (35–215) resulted in a phospho-signal at approximately 40 KDa that was absent in the GFP-ULK1 KD- and the GST-Lamin B1 coil 1 (35–215) S210A- containing reactions (Fig. 5c). In contrast, we could not detect a phospho-signal upon incubation with GST-Lamin B2 coil 1 (49–229) (Fig. 5c), indicating that ULK1 does not target Lamin B2 S224. Next, we tested whether the anti-Lamin B1 pS210 antibody cross-reacted with LB2 pS224 by Western blot. Indeed, a signal was visible in the reactions where AurA and PLK1 were incubated with the WT GST-Lamin B2 coil1 (49–229) but not with S224A (Fig. 5d). We assume that Lamin B2 phospho-S224 is not recognized in total cell lysates because we did not observe a signal in mitotic Lamin B1 KO cells by western blot (Fig. S2c). However, we cannot rule out that at least part of the signal we detected by immunofluorescence in our experiments was due to Lamin B2 S224 phosphorylation. Taken together, these data show that AurA and PLK1 can phosphorylate B-type lamins in the rod domain coil 1 *in vitro*, while ULK1 only targets Lamin B1.

**Figure 5:**
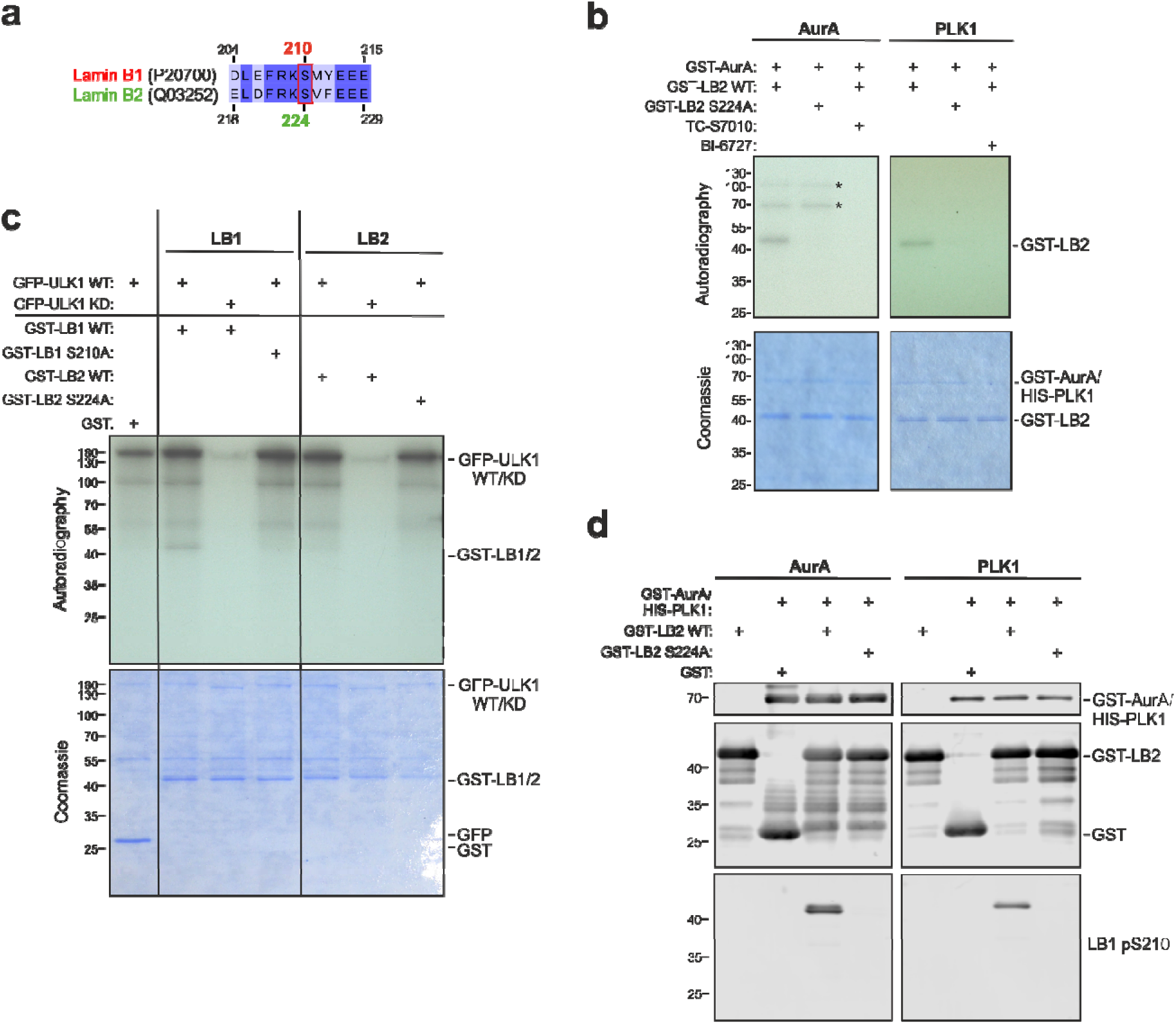
PLK1 and AurA can phosphorylate Lamin B2 at S224 *in vitro.* **a)** Protein sequence alignment of human Lamin B1 and Lamin B2 around the region containing S210 in Lamin B1 showing the presence of S224 in the corresponding position in Lamin B2. UniProt accession numbers are indicated in brackets. **b)** Radioactive *in vitro* kinase assays with purified GST-Lamin B2(49–229) WT or S224A and either GST-AurA (left panel) or HIS-PLK1 (right panel). The AurA inhibitor TC-S7010 or the PLK1 inhibitor BI-6727 (each used at 100 μM) were co-incubated with the kinase reactions as negative controls. After incubation with [ϒ-^32^P] ATP, the proteins were separated via SDS‒PAGE and exposed to film (autoradiography). Asterisks (*) in the AurA panel indicate GST-AurA auto-phosphorylation. **c)** Radioactive *in vitro* kinase assay with purified GFP-ULK1 WT or GFP-ULK1 KD (kinase-dead; D165A) and recombinant GST-Lamin B1 (35–215) (WT and S210A) or GST-Lamin B2 (49–229) (WT and S224A). After incubation with [ϒ-^32^P] ATP in the presence of kinase buffer, the samples were separated via SDS‒PAGE and exposed to film (autoradiography). **d)** Non-radioactive kinase assays with purified GST-Lamin B2 (49–229) WT or S224A and either GST-AurA (left panel) or HIS-PLK1 (right panel). After the kinase reaction, the samples were subjected to immunoblotting with the indicated antibodies. For all blots, LB1= Lamin B1, LB2 = Lamin B2.

### Lamin B1 S210 phosphorylation contributes to the maintenance of spindle integrity

We investigated the functional role of Lamin B1 S210 phosphorylation by establishing stable HeLa Lamin B1 KO cell lines expressing HA-Lamin B1 WT, phospho-abrogating S210A, or phospho-mimetic S210D variants. After confirming the similar expression levels of each variant and their proper targeting to the nuclear periphery (Fig. 6a, Fig. S5a), we first analyzed whether S210 phosphorylation is important for cell cycle progression but were unable to detect significant differences among the cell lines (Fig. S5b). Given the observed association of Lamin B1 pS210 with peri-centrosomal regions during prophase and with the spindle in later mitotic stages (Fig. 2c), we focused on analyzing spindle morphology. Co-staining of the cells with antibodies against the centrosomal protein ϒ-Tubulin and α-Tubulin by immunofluorescence revealed a substantial threefold increase in the number of mitotic cells displaying more than two centrosomes in the HA-Lamin B1 S210A variant compared to cells expressing HA-Lamin B1 WT or HA-Lamin B1 S210D (Fig. 6b). Most of the HA-Lamin B1 S210A cells with more than two centrosomes had three to four of them, leading in some cases to the formation of multipolar spindles (Fig. 6b, lower panel S210A). Because AurA and PLK1 activities have been linked to proper spindle orientation and function (*48, 49*), we also analyzed the angle and length of the spindle in metaphase cells that achieved bipolarity. We observed a slight decrease in the spindle length in HA-Lamin B1 S210A cells compared to that in WT and S210D cells (Fig. 6c). In contrast, no differences were detected in the spindle angles among these cell lines (Fig. S5c), suggesting that S210 phosphorylation contributes to only some aspects of mitotic spindle regulation. We additionally analyzed whether S210 phosphorylation is required for targeting HA-Lamin B1 to the spindle by staining each cell line with anti-HA antibody and scoring cells in metaphase. An enrichment of the signal at both sides of the metaphase plate, corresponding to the localization of the mitotic spindle, was observed in all three cell lines (Fig. 6d, red arrows). This indicates that phosphorylation of S210 alone is not sufficient for Lamin B1 targeting to the mitotic spindle. Finally, to analyze whether the observed anomalies in spindle structure delay mitotic progression, we performed live-cell imaging throughout mitosis in cells expressing either Lamin B1 WT or phospho-mutants. For this, we generated Lamin B1 KO cell lines expressing H2B-mCherry as a DNA marker and EGFP-Lamin B1 WT, S210A, or S210D. The similar expression levels of the GFP-Lamin B1 proteins and the identity of the cell lines were confirmed by immunoblotting with anti-LB1 and anti-LB1 pS210 antibodies in cells arrested in mitosis with paclitaxel (Fig. S6a). We synchronized the cell lines by double-thymidine block and stained microtubules with the live stain SPY650-Tubulin one hour before imaging. All three cell lines displayed proper targeting of GFP-Lamin B1 to the nuclear periphery at the beginning of the imaging, and cells progressed into mitosis synchronously (Fig. S6b). The mitotic duration was determined as the time elapsed from the moment of nuclear envelope breakdown (NEB) until anaphase onset (AO) (Fig. S6c). Various morphological and intensity features of each nucleus were used for the automated classification by a supervised machine learning module as mitotic (e.g, changes in DNA fluorescence intensity and nuclear area; a detailed analysis pipeline is presented in Table S5) and for their tracking until chromosome segregation occurred. Our results show a mild increase in the mitotic duration in cells expressing the S210A variant (44.43 ± 13.56 min) compared to the WT protein (42.77 ± 11.83 min), whereas S210D-expressing cells exhibited a slightly accelerated mitosis compared to WT cells (41.59 ± 10.56 min) and the difference between the two variants was significantly pronounced (Fig. 6e). These results suggest that modulation of Lamin B1 S210 phosphorylation during mitosis is important for timely mitotic progression, probably through the maintenance of proper spindle structure.

**Figure 6:**
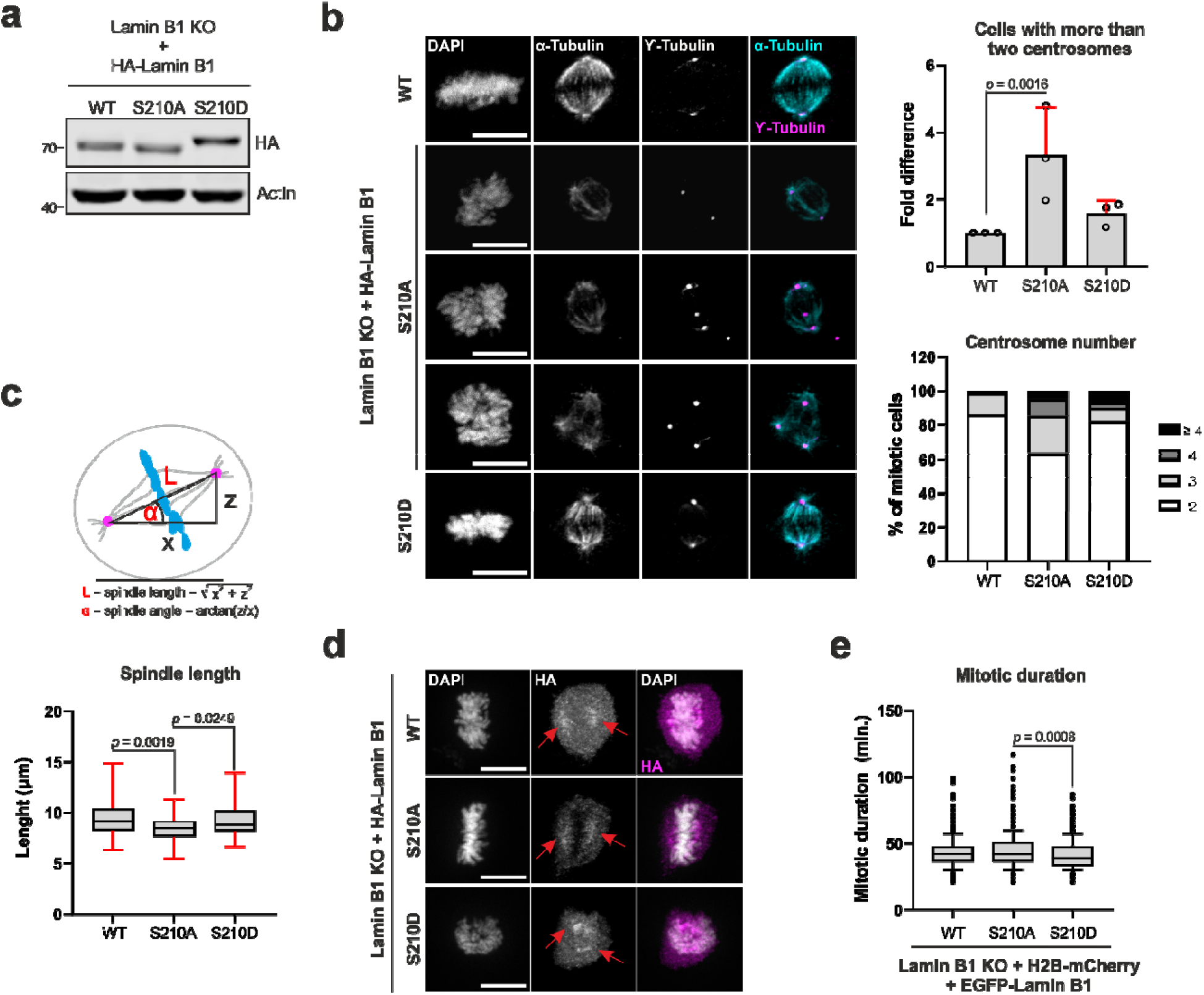
Lamin B1 S210 phosphorylation contributes to spindle integrity and mitotic progression. **a)** Lysates from HeLa Lamin B1 KO cells reconstituted with HA-Lamin B1 WT, S210A or S210D were processed for immunoblotting with the indicated antibodies. **b)** Representative images of reconstituted HeLa cells synchronized in mitosis by double-thymidine block and processed for immunofluorescence with ϒ-Tubulin and α-Tubulin antibodies. Cells in metaphase are shown. Scale bars, 10 μm. The graphs show the quantification of the proportion of mitotic reconstituted cells with more than two centrosomes relative to that of WT cells (upper graph) and the number of centrosomes in reconstituted mitotic cells (lower graph). The data in the upper graph are presented as the average + SD of three independent experiments. The total number of cells analyzed were: n_WT_ = 72, n_S210A_ = 82, n_S210D_ = 61 cells. The *p* value was determined by two-sided Fisher’s exact test. **c)** Upper panel: schematic representation of the mitotic spindle length and angle measurements from the data in b). x corresponds to the distance between centrosomes in projected images, and z is the number of z-stacks (1μm each) between both centrosomes. Lower panel: quantification of spindle length in each cell line. The data presented are pooled from three independent experiments and lines correspond to the minimum and maximum values. The total number of cells analyzed were: n_WT_ = 61, n_S210A_ = 51, n_S210D_ = 50 cells. The *p* values were calculated by one-way ANOVA with Tukey’s post hoc test. **d)** Representative images of reconstituted HeLa cells in metaphase processed for immunofluorescence with anti-HA antibody. Red arrows indicate the HA staining in a position corresponding to the mitotic spindle. Scale bars, 10 μm. **e)** Quantification of the mitotic duration in Lamin B1 KO cells expressing H2B-mCherry and GFP-Lamin B1 WT or phospho-mutants, from nuclear envelope breakdown until anaphase onset. Data shown are pooled from three independent experiments and lines correspond to the 10-90 percentile values. The total number of cells analyzed were: n_WT_ = 496, n_S210A_ = 501, n_S210D_ = 468 cells. The *p* value was calculated by one-way ANOVA with Tukey’s post hoc test.

### Lamin B1 phospho-S210 interacts with regulators of mitotic spindle function

To gain insight into the mechanism by which Lamin B1 pS210 contributes to spindle function, we used our specific phospho-antibody to immunopurify Lamin B1 pS210 from paclitaxel-arrested cells and subsequently detected co-purifying proteins by mass spectrometry (n=3). As a negative control, we used lysates treated with λ phosphatase prior to immunopurification and considered specific the hits that were significantly enriched (FDR<0.05) and that showed at least a two-fold enrichment in the experimental samples compared to the control. Based on these criteria, we obtained a list of 156 potential Lamin B1 pS210 interactors (Data S1), among which AurA was present (and PLK1 with a *p* value <0.05). Notably, other proteins involved in mitotic spindle assembly and the regulation of spindle size were found, such as NUMA1 (*50*), TPX2 (*51*), NUSAP1 (*52*) and the motor proteins KIFC1 (*53*) (HSET; also known as kinesin-14) and KIF5B (*54*) (Fig. 7a). We validated a subset of proteins by immunopurification and immunoblotting and confirmed that NUMA1 and PLK1 are relatively stable Lamin B1 pS210 interactors (Fig. 7b, lane 1). We also confirmed reported interactions between PLK1 and NUMA1 (*48*) (Fig. 7b, lanes 4 and 6) and between PLK1 and AurA (Fig. 7b, lanes 5 and 6). For the motor protein KIF5B, we detected a weak band in the Lamin B1 pS210, Lamin B1, and PLK1 immunopurifications, and in the KIFC1 pulldowns we detected a higher-migrating Lamin B1 band, which likely corresponds to modified Lamin B1 (Fig. 7b, lane 8). We were unable to detect Lamin B1 pS210 in the immunopurifications of all the other proteins (except for Lamin B1), suggesting that the abundance of co-purified endogenous Lamin B1 pS210 was below our detection limit by western blot analysis using the anti-Lamin B1 pS210 antibody (Fig. 7b, LB1 pS210 panel). We took particular interest in NUMA1 and KIFC1 as potential interactors of Lamin B1 pS210, given their known roles in maintaining spindle focusing (*55*) and centrosome clustering (*56*). To further validate these associations, we performed PLA analysis for Lamin B1 pS210 and NUMA1 or KIFC1 and detected a significant amount of signal in cells in prophase, pro-metaphase, and metaphase for both combinations compared to single-antibody controls (Fig. 7c). Our data show that Lamin B1 pS210 is associated with a network of proteins involved in spindle assembly and spindle pole focusing.

**Figure 7:**
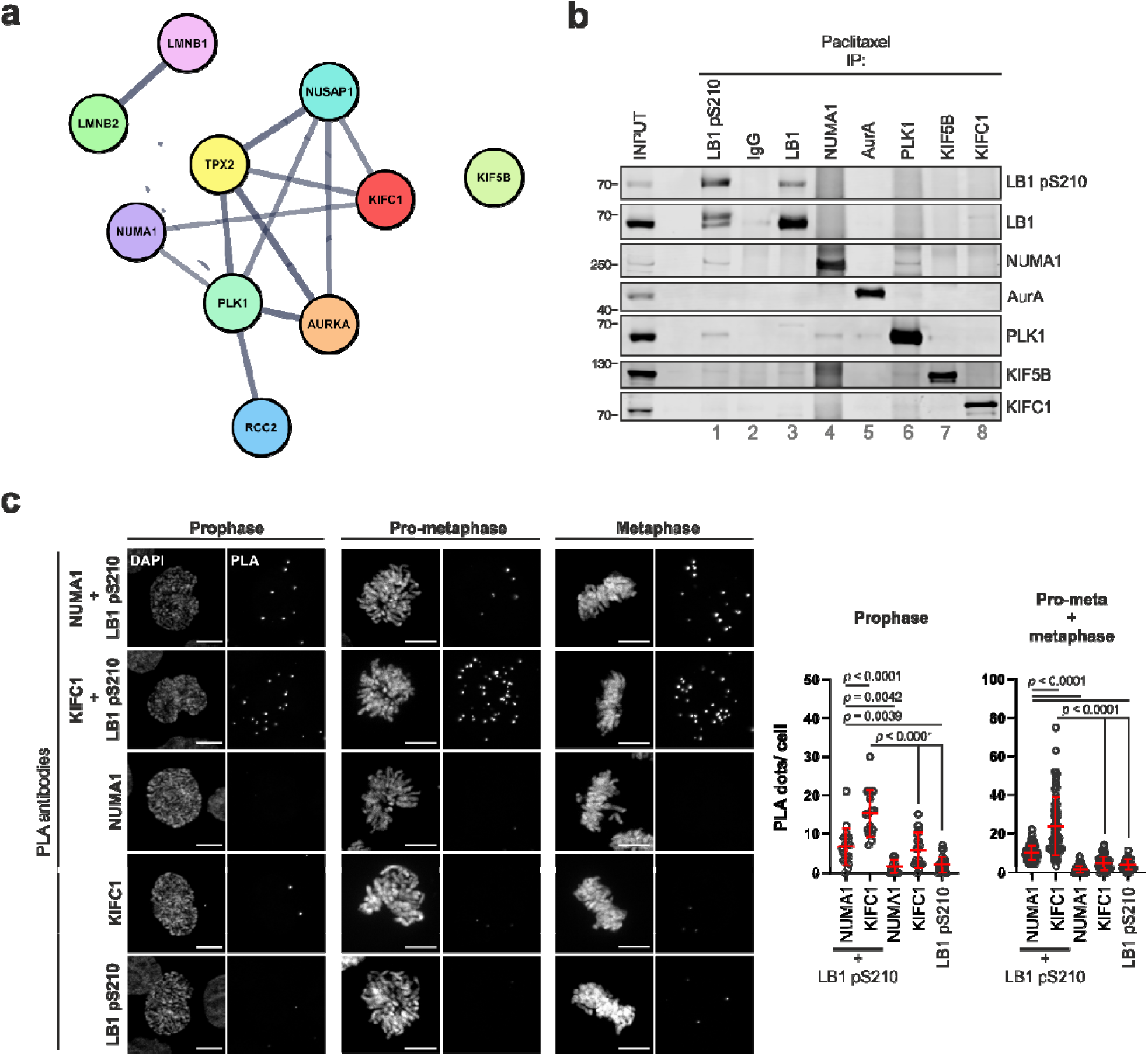
Lamin B1 interacts with a network of proteins involved in spindle assembly and spindle pole focusing. **a)** STRING network of a subset of proteins with known roles in spindle assembly and function that were detected by mass spectrometry co-purifying with Lamin B1 pS210 in paclitaxel-arrested cells. The full STRING network is displayed (edges indicate both functional and physical protein associations), where the width of the lines indicates the strength of data support. A medium confidence of 0.4 was used as the minimum required interaction score. **b)** HeLa cells were arrested with paclitaxel and equal amounts of total protein were immunopurified with the indicated antibodies. Co-purifying proteins were detected by immunoblotting with the antibodies depicted to the right of the panel. LB1 = Lamin B1.**c)** Representative images of PLA assays with the indicated antibodies in synchronized HeLa cells. Scale bars, 10 μm. The graphs show the quantification of PLA signals per individual mitotic cell. Pooled data from two independent experiments are shown, lines represent the means ± SDs. n_LB1 pS210/NUMA1_Prophase_= 18, n_LB1 pS210/KIFC1_Prophase_= 16, n_NUMA1_Prophase_= 15, n_KIFC1_Prophase_= 16, n_LB1 pS210_Prophase_= 25, n_LB1 pS210/NUMA1_Pro-meta/metaphase_= 76, n_LB1 pS210/KIFC1_ Pro-meta/metaphase_= 82, n_NUMA1_ Pro-meta/metaphase_= 85, n_KIFC1_ Pro-meta/metaphase_= 55, n_LB1 pS210_ Pro-meta/metaphase_= 64 cells. *P* values were determined by one-way ANOVA with Tukey’s post hoc test.

## Discussion

Phosphorylation events at the head and tail domains of lamins have been extensively characterized for their role in the disassembly of the lamin filament during mitosis (*21, 22, 57, 58*). Several phosphorylation events have been detected in the coiled-coil rod domain by high-throughput experiments, but, to our knowledge, it remains unclear whether these modifications affect lamin solubility or act as signaling platforms that mediate interactions with proteins involved in specific functions.

Here, we report that S210 in Lamin B1 and S224 in Lamin B2 are phosphorylated *in vitro* by the mitotic kinases AurA and PLK1, whereas only Lamin B1 can be further targeted by the autophagy-related kinase ULK1. By focusing on the analysis of Lamin B1 S210 phosphorylation with a specific antibody, we further demonstrated that Lamin B1 pS210 is a physiological modification that strongly increases during mitosis, initially arising at peri-centrosomal regions during prophase and subsequently localizing at the microtubule spindle in later mitotic stages. A spindle-specific localization of phospho-lamins was not apparent in previous studies using phospho-antibodies to visualize lamin localization during mitosis (*59*), which suggests that S210 phosphorylation, either alone or in concert with other modifications in the rod domain, may be specific for a spindle-related function. Particularly interesting is the fact that S210 seems to become phosphorylated before nuclear envelope breakdown and remain associated with microtubules until after nuclear envelope re-formation, arguing that this modification may be uncoupled from the broad CDK1- and PCK- dependent Lamin B1 phosphorylation that drives the disassembly of the nuclear lamina (*22*).

ULK1 plays an established role in initiating autophagy in response to cellular stress downstream of mTORC1. While most research has focused on its autophagic function, ULK1 has been widely demonstrated to play a plethora of roles in other processes. A first hint for a mitotic autophagy-independent role for ULK1 was reported in a study demonstrating that ULK1 can directly phosphorylate the spindle assembly checkpoint protein Mad1 and thus regulate mitotic exit (*29*). Our work identifies S210 in Lamin B1 as a novel mitotic ULK1 substrate by *in vitro* and proximity assays. However, it is likely that ULK1 plays a supporting role in the regulation of this modification, since the activities of the mitotic kinases AurA and PLK1 are also required for efficient Lamin B1 phosphorylation at S210. We envision a scenario in which all three kinases regulate S210 phosphorylation locally at distinct regions and mitotic stages. Indeed, it has been reported that both AurA and PLK1 localize to centrosomes during prophase, but in metaphase, AurA concentrates at the spindle poles, whereas PLK1 remains more mobile, transitioning between spindle poles and kinetochores (*60*). We detected ULK1 around the assembling microtubules during prophase and diffusing throughout the cytosol in metaphase. Thus, it is possible that all kinases have access to Lamin B1 at mitotic entry in a region proximal to the centrosomes, but that their individual contributions vary in later mitotic stages. Interestingly, we observed that pS210 levels are differentially affected by microtubule polymerization state, indicating the existence of a pool of Lamin B1 phosphorylated at microtubules and another that is independent of this scaffold. Although dissecting the specific involvement of each kinase is complicated due to their redundancy towards S210 phosphorylation and the extensive cross-regulation between AurA and PLK1 (*45*), it is possible that AurA reinforces and maintains a high S210 phosphorylation at the spindle poles during metaphase, whereas PLK1 and ULK1 are involved in keeping a more soluble pool of Lamin B1 pS210.

Our functional analyses of reconstituted cells expressing Lamin B1 WT, S210A, or S210D revealed that cell lines expressing the phospho-abrogating S210A variant exhibit a greater number of mitotic cells with more than two centrosomes and slightly shorter spindles than WT cells. Extra centrosomes are frequent in cancer cells and can be mechanistically linked to centrosomal amplification due to defects in centrosome duplication, prolonged G2 arrest, cytokinesis failure, and increased centriole length (*61*). Interestingly, mutation or depletion of several microtubule-regulating and mitotic motor proteins leads to the formation of multipolar spindles (*53, 54, 62, 63*). Multipolarity has also been linked to prolonged mitotic progression because altered spindle geometry leads to an abnormal attachment of kinetochores to spindle microtubules, activating the spindle assembly checkpoint (SAC) (*64*) and delaying the onset of anaphase. In accordance with the increased centrosome number in S210A-expressing cells, we also observed a moderate increase in mitotic duration compared to WT-expressing cells, and slightly accelerated progression in S210D-expressing cells. Thus, it is possible that the increased mitotic duration in S210A-expressing cells is related to checkpoint activation triggered by the observed extra-centrosomes. Conversely, a constitutive phosphorylation mimicked by the S210D variant would stabilize the spindle and prevent the formation of multipolar structures, allowing for a faster silencing of the SAC. Based on these observations, we propose that S210 phosphorylation contributes to proper mitotic progression by promoting spindle stability and bipolar spindle formation. In line with this idea, we identified NUMA1 and KIFC1, two key players in spindle pole focusing, as Lamin B1 pS210-interacting proteins. Interestingly, NUMA1 is also regulated by AurA-dependent phosphorylation on S1969, and abrogation of this modification leads to a reduction in spindle length (*65*) and an increase in the number of cells with multipolar spindles (*63*). These phenotypes resemble our observations in Lamin B1 S210A-expressing cells, suggesting that NUMA1 and Lamin B1 phosphorylation serve similar functions. The minus-end-directed motor KIFC1 contributes to spindle focusing by microtubule cross-linking, sliding, and bundling, and has attracted attention for its ability to induce clustering of supernumerary centrosomes, thereby promoting the survival of certain cancer types (*66–68*). It is possible that Lamin B1 pS210 supports the function of these specialized proteins by providing a binding scaffold at the mitotic spindle poles.

Overall, we propose that Lamin B1 belongs to a network of proteins regulated by upstream kinases such as ULK1, AurA, and PLK1, which control spindle focusing and length, ultimately impacting the duration of mitotic progression. Although it seems that S210 phosphorylation does not mediate Lamin B1 localization at the mitotic spindle (or matrix) by itself, it cannot be ruled out that it acts in concert with other modifications by enhancing Lamin B1 solubility or by mediating its interaction with other members of the network.

## Materials and Methods

### Cell lines and treatments

HeLa Lamin B1 KO cells were purchased from Abcam (# ab255404). 293T cells were purchased from ATCC (Cat# CRL-3216). Flp-In™ T-Rex™ 293 cells inducibly expressing GFP, GFP-ULK1 WT or GFP-ULK1 KD were generated previously (*40*). HeLa, 293T and Flp-In™ T-Rex™ 293 cells were cultured in Dulbecco’s Modified Eagle Medium (DMEM; Thermo Fisher Scientific, #41965039), and HCT116 cells were cultured in McCoy’s 5A (1X) + GlutaMAX™-I (Thermo Fisher Scientific #36600-021). Both media were supplemented with 10% fetal bovine serum (FBS; Sigma-Aldrich, #F0804), 4.5 g/l D-glucose, 100 U/mL penicillin and 100 µg/mL streptomycin (Thermo Fisher Scientific, #15140122). All cells were cultivated and treated at 37 °C and 5% CO_2_ in a humidified atmosphere. 293 cells were transfected using Lipofectamine® 2000 Transfection Reagent (Thermo Fisher Scientific #11668019). For synchronization experiments, HeLa cells were incubated in 9 μM RO-3306 (Sigma-Aldrich #SML0569) for 20 hr, washed twice in PBS and released into fresh media for the indicated times. For double-thymidine block, cells were incubated in 2 mM thymidine (Sigma-Aldrich #T9250) for 18 hr, washed 3 times in PBS, released into fresh media for 6 hr and re-blocked with 2 mM thymidine for 18 hr. After the second block, the cells were washed 3 times in PBS and released into fresh media for the indicated times. When kinase inhibitors were used, they were added 30 min before harvesting. Inhibitors for Aurora A (MLN8237, Biorbyt #orb154640), Aurora B (AZD1152-HQPA, Sigma-Aldrich #SML0268) and PLK1 (BI-6727; Cayman Chemicals #18193) were used at a final concentration of 500 nM, and ULK1 inhibitor (ULK101; Selleckchem #S8793) was used at 10 μM. For mitotic arrest HeLa or HCT116 cells were treated with 100 nM paclitaxel (Sigma-Aldrich #T7402) or 100 ng/ml nocodazole (Sigma-Aldrich #M1404) for 18 hr. Flp-In T-Rex 293 cells inducibly expressing GFP, GFP-ULK1 or GFP-ULK1-KD (Loffler *et al*., 2011) were induced with 0.1 μg/ml doxycycline for 18 hr.

### Antibodies

Antibodies against β-actin (WB: 1:20.000; Sigma-Aldrich #A5316), α-Tubulin (IF 1:500; Sigma-Aldrich #T5168; IF (for PLA) 1:100 Sicgen AB0134-200), GST (WB 1:2.000; Sigma-Aldrich #G7781), GFP (WB 1:2.000; ChromoTek #3H9), HIS tag (WB 1:2.000; Millipore #70796), GAPDH (WB 1:5.000; Abcam #ab8245 or WB 1:20.000; Cell Signaling Technologies # 5174 (Fig. S2a)), ULK1 (WB 1:1.000; Cell Signaling Technologies #8054 (Fig. 3a), or WB 1:2.000; Cell Signaling Technologies #4776, PLA 1:100 Santa Cruz Biotechnology # sc-33182); Lamin B1 (IF, PLA 1:200; Sigma-Aldrich # AMAB91251, WB 1:1.000, IP 1:500; Proteintech #12987-1-AP); Lamin B1 phospho-S210 (WB 1:500, IF 1:50, IP 1:150; developed in collaboration with Cell Signaling Technologies); Aurora A (WB 1:1.000; PLA 1:200 Cell Signaling Technologies #14475; IF 1:500, PLA 1:200 Cell Signaling Technologies #12100), PLK1 (WB 1:1.000, PLA 1:200, IP 1:1.000 Abcam #ab189139; IF 1:100, PLA 1:100 Santa Cruz Biotechnology # sc-17783), HA tag (WB 1:1.000, IF 1:500 Biolgend #901501), Flag tag (PLA 1:200, Sigma # F3165), ϒ-Tubulin (IF 1:200 Abcam #ab11317), Myc tag (WB 1:1.000 Cell Signaling Technologies #2278), H3 phospho-S10 (WB 1:1.000 Cell Signaling Technologies #3377), NUMA1 (WB 1:1.000, IP 1:250 Thermo Fischer Scientific # PA1-32451; PLA 1:100 Santa Cruz Biotechnology #sc-365532); KIFC1 (WB 1:500, IP 1:450 Proteintech #20790-1-AP; PLA 1:100 Santa Cruz Biotechnology #sc-100947), KIF5B (WB 1:1.000, IP 1:400 Proteintech #21632-1-AP) were used.

### Cloning

pGEX-Lamin B1 full-length, coil 1 (35–215) and coil 2 (216–386) were generated by PCR amplification from a Lamin B1 sequence optimized for bacterial expression (pET-20b(+)_HIS-LaminB1opt, Genscript) and cloned into pGEX-5X-3. pGEX-Lamin B1 head (1–34) and tail (387–586) were generated by back-to-back amplification of the full-length pGEX-Lamin B1. For site-directed serine/threonine-to-alanine mutagenesis of all pGEX-Lamin B1 constructs, the Q5 Site-Directed Mutagenesis Kit (New England Biolabs # E0554S) was used. pGEX-Lamin B2 coil 1 (49–229) was generated by PCR amplification of pCMV6-Lamin B2 (Origene # RC200807) and cloned into pGEX-5X-3 using Gibson Assembly Master Mix (New England Biolabs # E2611). For generation of pGEX-Lamin B2 coil 1 (49–229) S224A, the whole plasmid was amplified by PCR with back-to-back primers and subsequently phosphorylated and ligated with KLD Enzyme Mix (NEB #M0554S). Lamin B1 cDNA for expression in reconstituted HeLa Lamin B1 KO cells was obtained from the Dendra2-Lamin B1-10 plasmid (a gift from Michael Davidson, Addgene plasmid # 57728; http://n2t.net/addgene:57728; RRID:Addgene_57728). Lamin B1 cDNA was initially cloned into the pMSCVpuro-RFP-GFP vector by sequence-and-ligation-independent cloning (SLIC), in which single-stranded DNA was generated from both products by incubation with Klenow fragment and subsequent annealing in the presence of ligase buffer. Site-directed mutagenesis of this construct was performed to obtain pMSCVpuro-RFP-GFP-Lamin B1 WT, S210A and S210D. pMSCVpuro-HA-Lamin B1 WT, S210A and S210D were generated by replacing the RFP-GFP tag in each corresponding construct by back-to-back PCR amplification of the vector with a forward primer encoding the HA sequence, subsequent phosphorylation and ligation with KLD Enzyme Mix. All HA-Lamin B1 constructs were then sub-cloned into pMSCVhygro. Human ULK1 cDNA was cloned from p2CL21-flag-myc-ULK1 into pMSCVpuro by SLIC. pMSCVblast_GFP-Lamin B1 WT, S210A and S210D were generated by back-to-back amplification of pMSCVblast-RFP-GFP-Lamin B1 WT, S210A or S210D to remove the RFP sequence. pMSCVhygro_H2B-mCherry was generated by amplification of H2B cDNA (Histone H2B type 1-J was amplified) from total RNA from Jurkat cells and subsequent Gibson assembly with mCherry and pMSCVhygro amplified with overlapping overhangs. All the constructs and primers used in this study are listed in Table S2. For CRISPR/Cas9 editing, constructs containing ULK1 and ULK2 KO guide RNAs were generated using the system developed by the Zhang laboratory (*69*). For ULK1, the guide RNA used to target exon 1 was previously described (*29*). For ULK2, a guide RNA targeting exon 5 was generated by hybridization of DNA oligos (5’- CACCGACCTCGCAGATTATTTGCA -3’, 5’- AAACTGCAAATAATCTGCGAGGTC- 3’) and cloned into the Bbsl restriction site of the pSpCas9(BB)-2A-GFP (PX458) vector (a gift from Feng Zhang; Addgene plasmid #48138; http://n2t.net/addgene:48138; RRID:Addgene_48138).

### Retroviral transduction of HeLa cells

A total of 1.9 μg of the corresponding pMSCV retroviral plasmid was co-transfected with pCMV-VSVG into Plat-E cells using FuGENE 6 Transfection Reagent (Promega, #E2692) to produce pseudotyped recombinant retroviruses. After 48 hr, the retroviral supernatant was collected, and added together with 3 µg/ml polybrene (Sigma-Aldrich, #9268) to the target cells. After 72 hr, the cells were selected in medium supplemented with hygromycin (500 µg/ml; InvivoGen #ant-hg-1), blasticidin (100 ug/ml; InvivoGen # ant-bl-1) or puromycin (1 µg/ml; InvivoGen #ant-pr-1).

### Fractionation

HeLa cells at 60-80% confluency were trypsinized, washed once in PBS and re-suspended in extraction buffer A (20 mM Tris-HCl pH 7.6; 0.1 mM EDTA; 2 mM MgCl-6H_2_O) supplemented with protease and phosphatase inhibitor cocktail. After a 2-minute incubation at room temperature, the cells were transferred to ice and further incubated for 10 min. NP-40 was added to a final concentration of 1%, and the cells were homogenized by pipetting up and down. After centrifugation, the supernatant was collected as the cytoplasmic extract, and the nuclear pellet was washed twice in extraction buffer A supplemented with 1% NP-40. The washed nuclei were lysed in RIPA buffer containing protease and phosphatase inhibitors for 30 min on ice and then centrifuged, after which the supernatant was collected as the nuclear extract.

### Protein expression and purification

pGEX-Lamin B1/2 constructs were transformed into BL21 competent *E. coli.* Protein induction was performed by the addition of 1 mM IPTG to an exponentially growing culture for 4 hr at 23°C. Bacteria were pelleted and lysed by sonication in lysis buffer (50 mM Tris-HCl (pH 7.5), 300 mM NaCl, 5 mM EDTA, 5 mM EGTA, and 0.01% Igepal) supplemented with protease inhibitor cocktail, 50 mg/ml lysozyme and 20 U/ml DNAseI (Sigma-Aldrich # D4527). After centrifugation at 10,000 × g for 30 min, the lysate was incubated with glutathione sepharose 4B (Cytiva #17075605) for 1.5 hr at 4°C and washed 3 times with lysis buffer. Proteins were eluted from the beads by incubation in elution buffer (50 mM Tris base, 10 mM reduced L-glutathione, pH 8.8) for 30 min at 4°C with rotation.

### Immunoblotting, immunopurification and λ phosphatase treatment

Cells were harvested by scraping on ice, pelletized and flash-frozen until processing. For immunoblotting, cell pellets were lysed in RIPA buffer (50 mM Tris-HCl pH 8.0; 150 mM NaCl; 1% Triton X-100; 0.5% sodium deoxycholate; 0.1% SDS) supplemented with protease inhibitor cocktail (Sigma-Aldrich, #P2714) and phosphatase inhibitor cocktail (PhosSTOP; Roche # 04906837001) for 30 min on ice. Lysates were clarified by centrifugation at 18,000 rcf and 4°C for 20 min. Equal amounts of protein were separated via SDS‒PAGE and subsequently transferred to PVDF membranes (Merck Millipore, #IPFL00010). Membranes were blocked in 5% non-fat milk for 1 hr at room temperature, except when probed with anti-Lamin B1 phospho S210 antibody, for which the membranes were blocked in 5% BSA. Membranes were incubated with the indicated antibodies overnight at 4°C with rotation, washed three times with TBS-T, and probed with the appropriate IRDye-conjugated secondary antibodies for 1 hr at room temperature. After three final washes in TBS-T, the membranes were scanned in an Odyssey Infrared Imaging System (LI-COR Biosciences) and quantified using Image Studio Lite 4.0 (LI-COR Biosciences). For Fig. S2a, western blotting was performed as suggested by Cell Signaling Technology https://www.cellsignal.com/learn-and-support/protocols/protocol-western and detected with LumiGlo® (Cell Signaling Technologies #7003). For immunopurification, cell pellets were lysed in lysis buffer (20 mM Tris-HCl (pH 7.5), 137 mM NaCl, 1% Triton X-100, and 2 mM EDTA) supplemented with protease and phosphatase inhibitor cocktail for 30 min on ice. Lysates were clarified by centrifugation at 18,000 rcf and 4°C for 20 min and pre-cleared by incubating with Protein-A beads (Cytiva #17528001) for 4 hr at 4°C with rotation. Equal amounts of protein from the pre-cleared lysates were incubated with the indicated antibodies overnight at 4°C with rotation. Protein-A beads were added to each reaction tube and incubated for 4 hr at 4°C with rotation. Beads were washed three times in cold lysis buffer, and the samples were separated via SDS‒PAGE and immunoblotted as described above. For affinity purification of GFP, GFP-ULK1 and GFP-ULK1 KD from Flp-In™ T-Rex™ 293 cells, pellets were lysed in lysis buffer (50 mM Tris-HCl pH 7.5; 150 mM NaCl; 1 mM EDTA; 0.3% CHAPS) supplemented with protease and phosphatase inhibitor cocktail for 30 min on ice. Lysates were clarified by centrifugation at 18,000 rcf and 4°C for 20 min, and equal amounts of protein were incubated with GFP-trap® beads (ChromoTek, #gta-200) overnight with rotation. Beads were washed twice in high-salt wash buffer (50 mM Tris-HCl, pH 7.5; 300 mM NaCl; 1 mM EDTA; 1 mM EGTA) and twice in 1x kinase buffer A (5 mM Tris-HCl, pH 7.5; 10 µM EGTA; 100 µM DTT; 7.5 mM Mg(CH_3_COO)_2_). The bead pellets were used for kinase assays. For phosphatase treatment of lysates, the samples were incubated with λPP (8 U/µl; NEB #P0753) in the presence of 1x PMP buffer and 1x MnCl_2_ for 45 minutes at 30 °C. After incubation, the samples were separated by SDS‒PAGE and immunoblotted as described.

### Immunofluorescence and image analysis

Cells were seeded on glass coverslips and treated as indicated. For fixation, the cells were washed once in PBS and incubated in 4% paraformaldehyde for 10 min at room temperature. After 3 washes in PBS, the fixed cells were permeabilized in 0.5% Triton X-100 for 10 min at room temperature, blocked with 3% BSA (Roth, #8076) for 1 hr at room temperature and incubated with primary antibodies diluted in 3% BSA overnight at 4°C. The samples were washed 3 times with PBS, incubated with secondary antibodies diluted in 3% BSA for 1 hr at room temperature, washed twice with PBS and stained with 0.5 µg/ml DAPI (Roth, #6335.1) for 5 min. After a final wash in PBS, the cells were embedded in ProLong Glass Antifade Mountant (Thermo Fisher Scientific, #P36980). For phosphatase treatment, the samples were fixed and permeabilized, incubated with λPP (10 U/µl; NEB #P0753) in the presence of 1x PMP buffer and 1x MnCl_2_ for 18 hr at 30 °C, and then processed for immunofluorescence as described above. Images were collected with an Axio Observer 7 fluorescence microscope (Carl Zeiss Microscopy) using a 40x/1,4 Oil DIC M27 Plan-Apochromat objective (Carl Zeiss Microscopy) and an ApoTome 2 (Carl Zeiss Microscopy). For quantification of fluorescence intensity, z-stacks were projected in SUM mode, and the intensities of individual mitotic cells were measured with ImageJ. For correlation analysis, z-stacks were projected, and images of individual mitotic cells were analyzed with the Coloc2 plugin of ImageJ after background subtraction.

### Proximity ligation assay (PLA) and image analysis

PLA was performed using the Duolink® *In Situ* Detection Reagents Far Red (Sigma-Aldrich #DUO92013) according to the manufacturer’s protocol. Briefly, cells were seeded on coverslips, treated as indicated and then fixed in 4% paraformaldehyde for 10 min at room temperature. After 3 washes in PBS, the fixed cells were permeabilized in 0.5% Triton X-100 for 10 min at room temperature, washed again 3 times in PBS and blocked in Blocking Solution (Sigma-Aldrich #DUO82007) for 1 hr at 37°C. The antibodies were diluted in antibody diluent (Sigma-Aldrich #DUO82008) and added to the samples overnight at 4°C. To simplify the categorization of cells in each mitotic stage a goat anti-Tubulin antibody was added together with the PLA antibodies. After washing twice in Wash Buffer A, the Duolink PLA Probes (1:5 Duolink *In Situ* PLA Probe Anti-Mouse MINUS (Sigma-Aldrich #DUO82004) and 1:5 Duolink *In Situ* PLA Probe Anti-Rabbit PLUS (Sigma-Aldrich #DUO82002)) were applied for 1 hr at 37°C. The secondary anti-goat antibody Alexa 488 was added together with the probes, diluted at 1:500. The samples were washed twice in Wash Buffer A and the Ligation Solution was added for 30 min at 37°C. The samples were subsequently washed again twice in Wash Buffer A and incubated with the Amplification Solution for 100 min at 37°C. All incubations at 37°C were performed in a humidified chamber. Following two final washes with 1x Wash Buffer B, the cells were briefly washed with 0.01x Wash Buffer B and mounted with mounting medium containing DAPI (Sigma-Aldrich #DUO82040). The samples were imaged as described for immunofluorescence.

For quantification of the PLA dots in Fig. 1b, image stacks were projected, nuclei were segmented, and PLA dots were counted with a CellProfiler™ pipeline (https://cellprofiler.org/) included in Table S3. Briefly, nuclei segmentation was performed using DAPI staining. The nuclei were dilated by 10 pixels to include signals close to the nuclear periphery, and PLA dots were counted in the segmented ROIs. For analysis of mitotic cells image stacks were projected and mitotic cells were manually scored, categorized into mitotic stages and isolated. PLA dots from each individual cell were counted using CellProfiler™ with a pipeline included in Table S4.

### siRNA

For transient knockdown of ULK1, HeLa cells were seeded in 6-well plates and transfected with 30 nM ON-TARGETplus Human ULK1 siRNA-SMARTpool (Dharmacon, #L-L-005049-00-0010) or ON-TARGETplus Non-targeting Control Pool (Dharmacon, #D-001810-10-20) in the presence of RNAiMAX (Thermo Fisher, # 13778075). The cells were analyzed 48 hr after transfection.

### Generation of CRISPR KO cells

Cells were -transfected with 2 µg of each pX458_ULK1 sgRNA and pX458_ULK2 sgRNA by electroporation using the Amaxa® Cell Line Nucleofector® Kit R (Lonza, #VCA-1001) according to the manufacturer’s instructions. Four days after transfection, the GFP-positive cells were sorted, and single-cell clones were generated. ULK1 knockout was validated by immunoblotting, and ULK2 knockout was validated by sequencing the ULK2 locus due to the lack of specific anti-ULK2 antibodies. For this purpose, genomic DNA was isolated using the GeneJET Genomic DNA Purification Kit (Thermo Fisher Scientific, #K0721), and the genomic locus of ULK2 around the guide RNA targeting site was amplified via PCR using the following primers: 5′- AGATAATTACATCATTGTGTTGGG-3′ and 5′-GAAGGAGAGGAATTCTATCGGG-3′. PCR products were cloned into the pCR™ 2.1-TOPO™ TA-vector using a TOPO™ TA Cloning™ Kit (Thermo Fisher Scientific, #450641), and the obtained sequences were aligned with the genomic ULK2 sequence (https://www.ncbi.nlm.nih.gov/gene/9706) since the editing site is located close to the exon 5/intron junction in the ULK2 gene. Two sequences were recovered—a T insertion and a T deletion three base pairs upstream from the PAM—leading to a frameshift and premature stop codon. A ULK1-negative clone harboring edited ULK2 was selected for the experiments.

### Radioactive *in vitro* kinase assays

For radioactive kinase assays, 500 ng of recombinantly expressed GST-Lamin B1 or GST-Lamin B2 fragments were incubated with 500 ng of GST-ULK1 (1–649) (Sigma-Aldrich #SRP5096), GST-Aurora A (Sigma-Aldrich #A1983) or HIS-PLK1 (Sigma-Aldrich #P0060). For kinase reactions with affinity-purified GFP-ULK1 WT or kinase-dead (KD), 500 ng of GST-Lamin B2 or GST-Lamin B1 fragments were incubated with each bead-bound kinase. For assays in Fig. 5b the Aurora A inhibitors TC-S7010 (Sigma-Aldrich #SML0882) and BI6727 were used at 100 μM. All the reactions were performed in the presence of 2 µM ATP, 10 µCi [ϒ-^32^P]-ATP (Hartmann Analytic), 2.5 mM Tris/HCl (pH 7.5), 5 µM EGTA, 50 µM DTT, and 3.75 mM Mg(CH_3_COO)_2_ for 45 min at 30°C and 450 rpm. After incubation, the samples were separated by SDS‒PAGE, gels were stained with Coomassie blue, and after drying, exposed to radiographic film in the dark.

### Non-radioactive *in vitro* kinase assays

To identify ULK1- or ULK3-dependent phosphorylation at residues in Lamin B1, an *in vitro* kinase assay with non-radioactive ATP was performed. For this purpose, 2 μg of purified full-length Lamin B1 (Origene #TP301604) was incubated with 1 μg of either GST-ULK1 (1–649) or HIS-ULK3 (Sigma-Aldrich #SRP5098) in the presence of 6 μM ATP and 2.5 mM Tris/HCl (pH 7.5), 5 µM EGTA, 50 µM DTT, and 3.75 mM Mg(CH_3_COO)_2_ for 1 hr at 30°C and 450 rpm. After incubation, the reactions were flash-frozen and processed for mass spectrometry. For non-radioactive kinase assays with GST-Lamin B1 or GST-Lamin B2, 500 ng of each fragment was incubated with 500 ng of either GST-Aurora A or HIS-PLK1. For assays with affinity-purified GFP-ULK1 WT and KD, bead-bound kinase was incubated with 500 ng of GST-Lamin B1 fragments. All the reactions were performed in the presence of 3 μM ATP, 2.5 mM Tris/HCl (pH 7.5), 5 µM EGTA, 50 µM DTT, and 3.75 mM Mg(CH_3_COO)_2_ for 1 hr at 30°C and 450 rpm. After incubation, the samples were subjected to immunoblotting with the indicated antibodies.

### Flow cytometry

To determine the cell cycle distribution, a BrdU uptake assay was performed. The cells were treated with 30 μM BrdU (Sigma #B9285) for 1 hr at 37°C, trypsinized, counted, aliquoted and fixed in cold 70% ethanol. Fixed cells were incubated in 2 N HCl with 0.5% Triton X-100 for 30 min at room temperature with rotation. After one wash in 0.1 M sodium tetraborate and one wash in washing buffer (1% BSA in PBS), the cells were re-suspended in 1:100 mouse anti-BrdU antibody (Santa Cruz Biotechnology # sc-32323) diluted in antibody buffer (0.5% Tween-20; 1% BSA in PBS) and left rotating overnight at 4°C. For determination of the mitotic index, cells were fixed in cold 70% ethanol, permeabilized in 0.25% Triton X-100 on ice for 15 min and incubated with a rabbit anti-H3 phospho-S10 antibody (Cell Signaling Technology #D2C8) diluted 1:1600 in wash buffer overnight at 4°C. All the samples were washed twice in wash buffer and incubated with 1:500 anti-mouse (Invitrogen #A32766) or anti-rabbit Alexa 488 (Invitrogen #A32731) diluted in antibody buffer for 30 min at room temperature. After a final wash in washing buffer, each cell pellet was re-suspended in 0.3 ml of PBS containing 3 μM Draq7 to stain the DNA (Abcam #ab109202), filtered through a cell strainer, processed in a BD LSR Fortessa flow cytometer with FACSDIVA software and analyzed with FlowJo software.

### Live cell imaging

HeLa Lamin B1 KO cells expressing H2B-mCherry and GFP-Lamin B1 WT, S210A or S210D were seeded in triplicates on 96-well Phenoplates (Revvity #6055302) at a density of 6.500 cells/well and synchronized by double-thymidine block as described above. The day of the experiment the cells were released form the second block and stained with SPY650-Tubulin (Spirochrome; 1:1000 dilution in cell culture media) for 1 hour before imaging (7 hours after release) at 37 °C and 5 % CO_2_. After incubation, the cells were placed into the preheated microscope to warm up again. Images were recorded at an Operetta CLS high-content screening system (Revvity) with environmental control (37 °C, 5 % CO_2_). Images were acquired with software Harmony 5.2 (Revvity, 2024) and saved as tiff-files. Time series images were captured every three minutes with a Zeiss EC Plan-Neofluar 40x air objective (NA 0.75) in widefield mode for at least three hours. For each cell line, three wells were imaged, with eight unconnected fields captured per well. Three channels were captured for evaluation of mitotic duration with three z-planes each. For excitation a 475 nm LED was used for EGFP (em. filter 500-550 nm), a 550 nm LED for mCherry (em. filter 570-650 nm) and a 630 nm LED for SPY650-Tubulin (em. filter 665-760 nm). The procedure was performed on three individual days, leading to three biological replicates from different cell passages for each cell line.

### Quantification of mitotic duration

Images were analyzed using Harmony 5.2 image analysis module (Revvity, 2024) and processed as flatfield-corrected maximum intensity projections. First, nuclei were detected as individual objects based on mCherry fluorescence and tracked over time. Nuclei touching image borders were omitted. Various morphological, intensity and texture features were measured for each nucleus to use the supervised machine learning module (PhenoLOGIC™) for classification of nuclei into mitotic or non-mitotic. Track and object data was exported from Harmony and processed with KNIME 5.3 (KNIME AG) (*70*). This processing included removal of debris and segmentation artefacts. For quantification only nuclei were considered that were present in the first image of each time series, not in mitosis at that time and underwent a cell division within the observation period of three hours. Cell cycle duration was quantified as time from entering mitotic class until splitting recognized by the tracking module. In total at least 468 nuclei were evaluated per cell line.

### Mass spectrometry analyses

Samples of in vitro kinase assays were digested in solution using trypsin and phosphopeptides were purified as described previously (*71*). For interactome studies, samples were digested on-beads with trypsin following the FASP protocol (*72*). In all cases, peptides were purified by STAGE tips prior LC-MS/MS measurements. The LC-MS/MS measurements were performed on a QExactive HF-X mass spectrometer coupled to an EasyLC 1000 nanoflow-HPLC. The mass spectrometer was operated in data-dependent mode for in vitro kinase assays as described previously (*71*) and in data-independent mode for interactome studies. Briefly, after each survey scan (mass range m/z = 350 – 1,200; resolution: 120,000) 28 DIA scans with an isolation width of 31.4 m/z were performed covering a total range of precursors from 350-1,200 m/z. AGC target value was set to 3 x 10^6^, resolution to 30,000 and normalized collision energy to 27%. Data were analyzed using Spectronaut software version 18.5 (Biognosys) with standard settings (without imputation) in direct DIA mode using reference proteome of human (UniProt, full length) and common contaminants. Further data processing and statistical analyses were done using Perseus (*73*). Statistical significance was calculated using a one-sided unpaired student T-test with FDR corrected p values <0.05.

### Statistical analysis

For densitometry quantification of the Lamin B1 phospho-S210 bands, the density of each protein band was divided by the average density of all the bands on the membrane. The same procedure was used for the total Lamin B1 bands. The resulting values from Lamin B1 phospho-S10 were normalized to the corresponding total Lamin B1 values. Fold changes were calculated by dividing each normalized density ratio by the average of the density ratios of the indicated control lane. For comparisons between conditions in one group, one-way ANOVA was used; for comparisons between groups, two-way ANOVA was performed. For analysis of categorical data, Fisher’s exact T test was used. All analyses were performed using GraphPad Prism 8 (GraphPad Software).

## Acknowledgments

We would like to thank members of the Stork laboratory and Dr. Lea Esser for sharing reagents, providing technical support and revising the manuscript. We thank Carole Roubaty and Michael Stumpe for mass spectrometry support. We thank Feng Zhang and Michael Davidson for providing plasmids (Addgene plasmid #48138 and Addgene plasmid # 57728, respectively).

## Funding

Human Frontier Science Program (HFSP) Post-doctoral Fellowship for MJM (LT 000993/2013).

Deutsche Forschungsgemeinschaft (DFG)- STO 864/4-3 (project #267192581).

Deutsche Forschungsgemeinschaft (DFG)- STO 864/9-1 (project #542770124).

Deutsche Forschungsgemeinschaft (DFG)- GRK 2158 (project #270650915).

Deutsche Forschungsgemeinschaft (DFG)- GRK 2578 (project #417677437).

Deutsche Forschungsgemeinschaft (DFG)- INST 208/760-1 FUGG for the Revvity Operetta CLS high-content screening automated microscope system.

## Author contributions

Conceptualization: MJM, BS

Methodology: MJM

Software: ABH

Investigation: MJM, CK, AS, NB, LB, YS, PN, ZH

Resources: GK, ZH, JD, ABH

Data Curation: MJM, ABH

Visualization: MJM

Supervision: MJM, JD, BS

Writing—original draft: MJM

Writing—review & editing: MJM, DJ, BS

Funding acquisition: BS

## Data and materials availability

The mass spectrometry proteomics data have been deposited in the ProteomeXchange Consortium (*74*) via the PRIDE (*75*) partner repository with the dataset identifier PXD055739. The live-cell imaging data will be deposited in the BioImage Archive (https://www.ebi.ac.uk/bioimage-archive/) after publication.

