## Supplementary Information for "Mitotic phosphorylation of Lamin B1 rod domain by ULK1 and Aurora A/PLK1 promotes spindle function"

María José Mendiburo *et al.*

* María José Mendiburo.

**This PDF file includes:**

Figs. S1 to S6

Tables S1 to S5

Data S1 caption

**Other Supplementary Materials for this manuscript include the following:**

Data S1

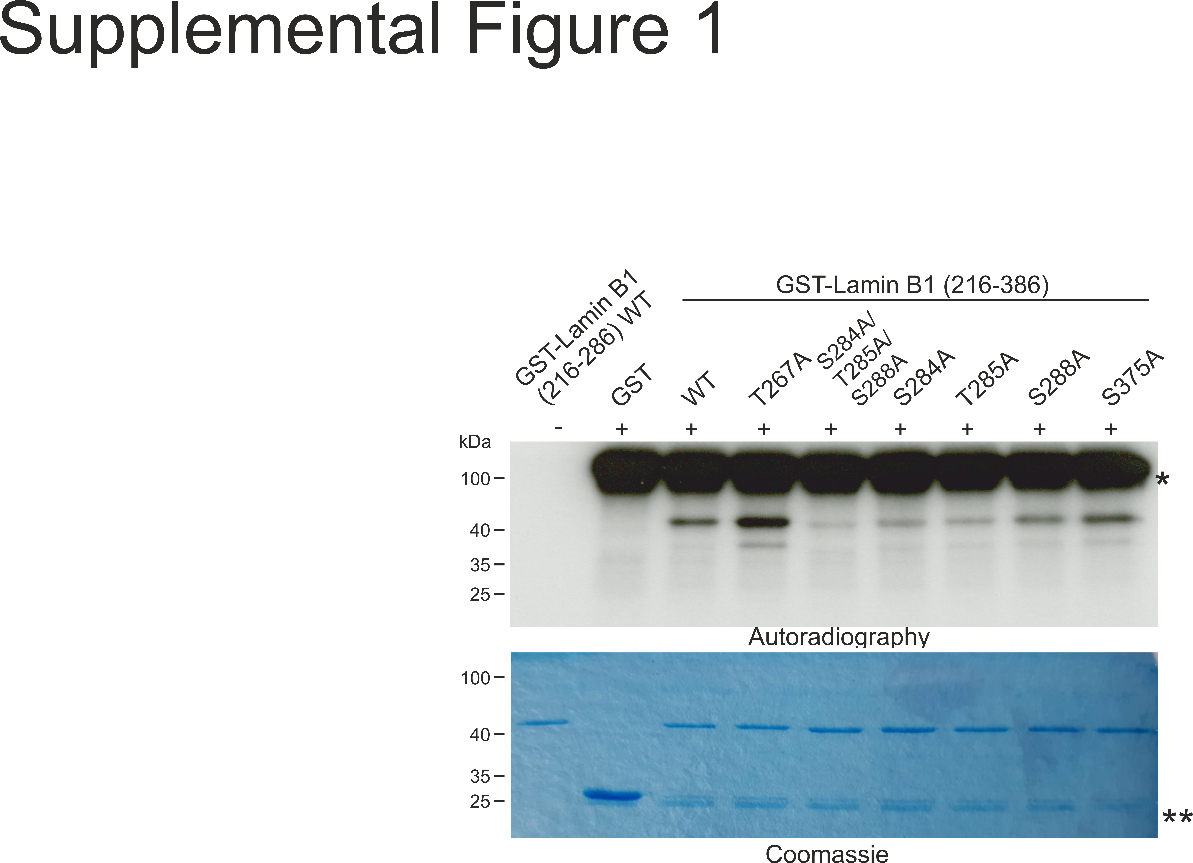

**Fig. S1: ULK1 phosphorylates Lamin B1 coil 2 within the rod domain**. Radioactive *in vitro* kinase assays with GST-ULK1(1-649) and recombinant GST-Lamin B1 coil 2 (216--386) fragment either WT or T267A, S284A, T285A, S288A and S375A variants. A triple S284A/T285A/S288A variant was also included due to the close proximity of the residues. After incubation with [ϒ-^32^P] ATP reactions were separated in SDS-PAGE gels and exposed to film (autoradiography). * indicates GST-ULK1 auto-phosphorylation, ** indicates free GST.

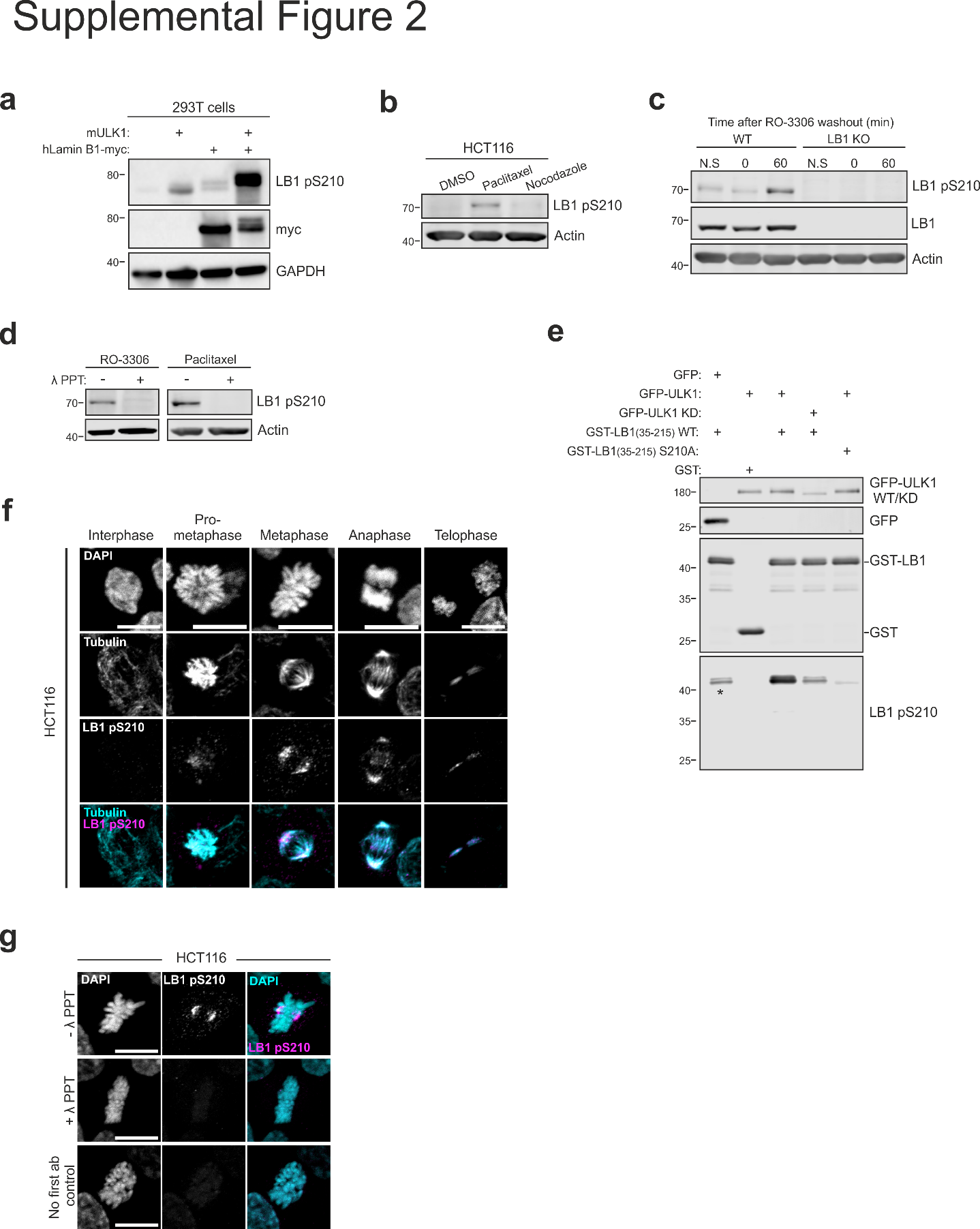

**Fig. S2: Validation of the anti-Lamin B1 S210 phospho-antibody**. **a)** 293T cells were transfected with mouse ULK1 and human Lamin B1-myc constructs individually or in combination. Lysates were immunoblotted with the indicated antibodies. LB1 = Lamin B1 for all panels. **b)** Lysates from HCT116 cells arrested with 100 nM paclitaxel or 100 ng/ml nocodazole for 18 hours were immunoblotted and probed with the indicated antibodies. **c)** Lysates from WT and Lamin B1 KO Hela cells synchronized in mitosis by RO-3306 treatment and release for 60 minutes were immunoblotted and probed with the indicated antibodies. N.S = non synchronized. **d)** Lysates from Hela cells synchronized in mitosis by arrest with RO-3306 and release for 60 minutes or arrested with 100 nM paclitaxel for 18 hours were incubated or not with λ-phosphatase (λ PPT). After immunoblotting samples were probed with the indicated antibodies. **e)** *In vitro* kinase assay with purified GFP-ULK1 WT or GFP-ULK1 KD (kinase-dead; D165A) and recombinant GST-Lamin B1 (35-215) fragment either WT or S210A variant. After kinase reaction samples were immunoblotted and detected with the indicated antibodies. Asterisks in the anti-Lamin B1 pS210 (LB1 pS210) blot indicates signals present in GST-Lamin B1(35-215) WT without kinase (incubated with GFP as control). **f)** Representative images (z-stack projections) of HCT116 cells fixed and processed for immunofluorescence with anti-Lamin B1 phospho S210 and anti- αTubulin antibodies. Scale bars, 10 μm. **g)** Representative images (z-stack projections) of HCT116 cells fixed, treated or not with λ-phosphatase (λ PPT) and processed for immunofluorescence with anti-Lamin B1 phospho S210 antibody. Scale bars, 10 μm**.**

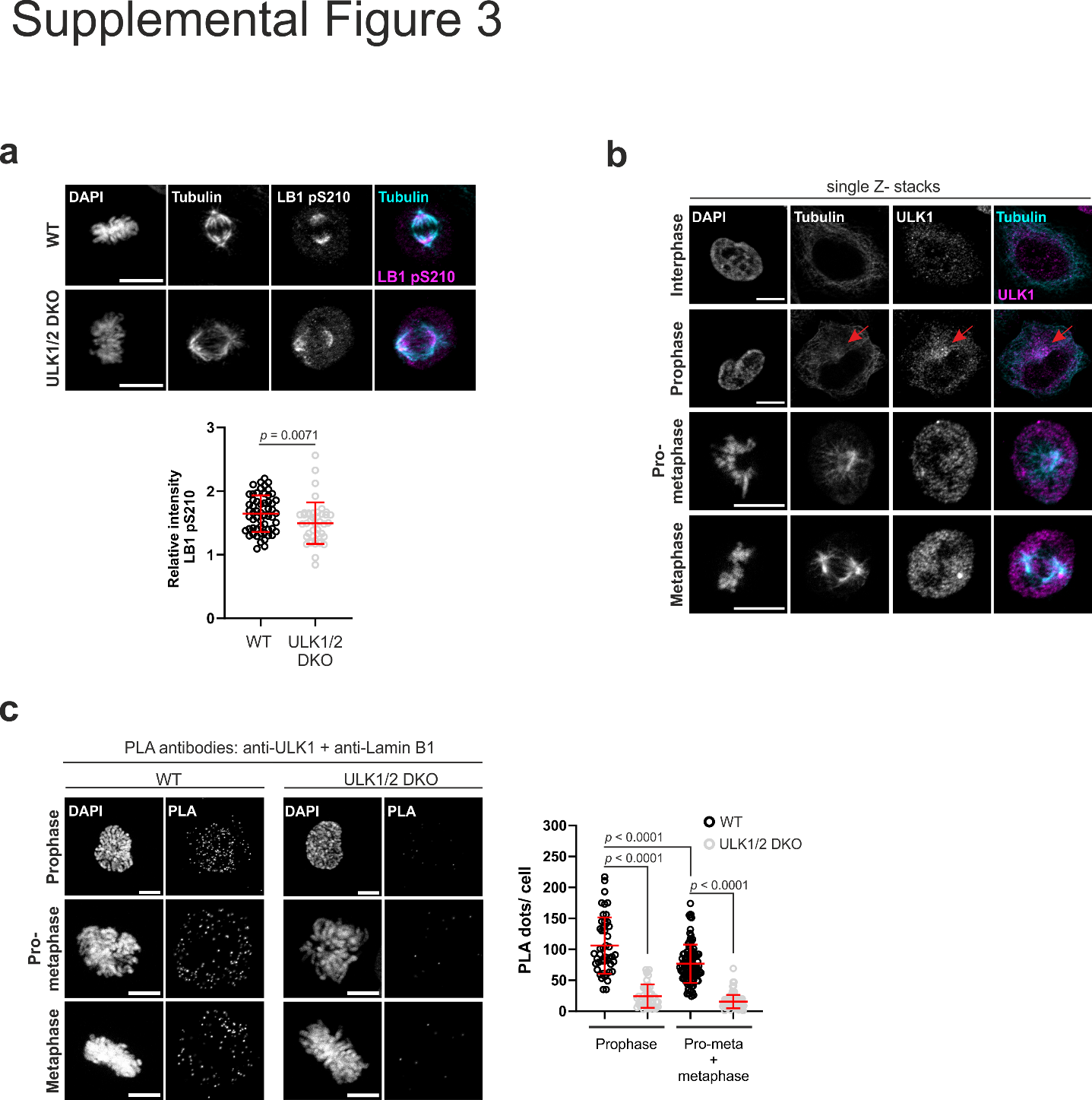

**Fig. S3: ULK1 contributes to Lamin B1 phosphorylation at S210**. **a)** Immunofluorescence images of HeLa WT or ULK1/2 DKO cells in metaphase stained with the indicated antibodies. Scale bars, 10 μm. Lower panel: The mean Lamin B1 pS210 fluorescence intensities were standardized to the average intensity of four interphase cells in the same image. The data shown are from two independent experiments, lines represent the means ± SDs. n_WT_ = 57 cells, n_ULK1/2 DKO_= 43 cells. The *P* value was determined with the Mann‒Whitney test. **b)** Representative images (single Z-stacks) of mitotic HeLa cells stained with anti-ULK1 and anti- tubulin antibodies. The red arrow indicates a peri-centrosomal region where ULK1 accumulates during prophase. Scale bars, 10 μm. **c)** PLA of WT and ULK1/2 DKO cells in mitosis using antibodies against ULK1 and Lamin B1. Scale bars, 10 μm. Right panel: Quantification of the number of PLA puncta in individual cells. The data were pooled from three independent experiments; the lines represent the means ± SDs. n_WT_Prophase_ = 48 cells, n_WT_Pro-meta/ metaphase_ = 101 cells, n_ULK1/2 DKO_Prophase_ = 43 cells, n_ULK1/2 DKO_Pro-meta/metaphase_= 93 cells. *P* values were determined by two-way ANOVA with Tukey´s post hoc test.

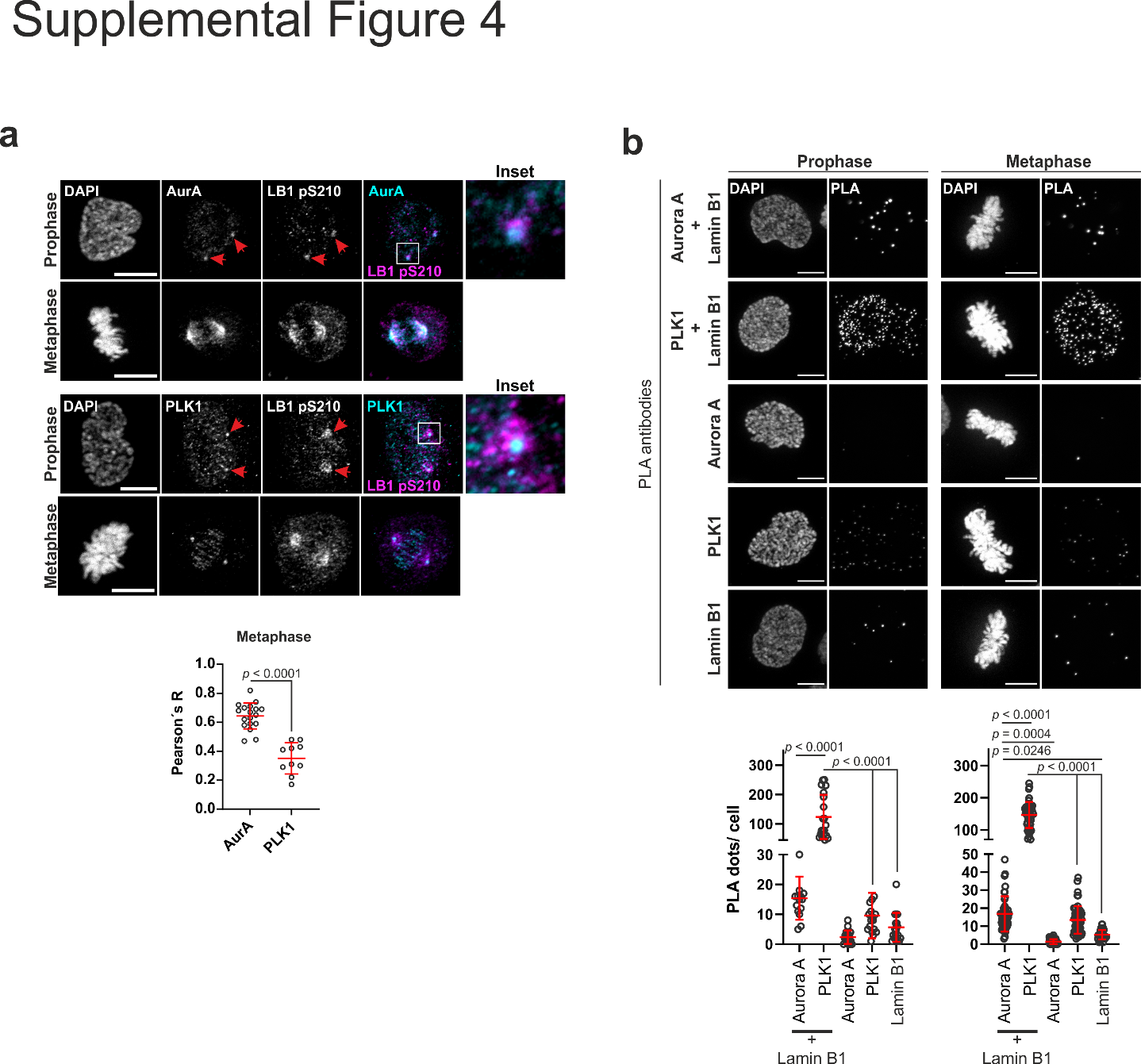

**Fig. S4: AurA and PLK1 also contribute to Lamin B1 phosphorylation at S210**. **a)** Representative images of mitotic HeLa cells stained with anti-Lamin B1 pS210 and anti- Aurora A (upper panel) or anti-PLK1 (lower panel) antibodies. The red arrowheads indicate Lamin B1 pS210 and each kinase in the centrosomal region during prophase. One centrosome of each staining pair is shown in the insets. Scale bars, 10 μm. The graph below shows the quantification of the correlation (Pearson´s R coefficient) between the Lamin B1 pS210 and AurA or PLK1 signals in metaphase cells like shown above. The data were pooled from two independent experiments; the lines represent the means ± SDs. n_AurA_ = 14 cells, n_PLK1_ = 10 cells. The *P* value was determined by the Mann‒Whitney test. **b)** Representative images of PLA assays with the indicated antibodies in mitotic HeLa cells. Scale bars, 10 μm. The graphs below show the quantification of PLA signals per individual mitotic cell. Pooled data from two independent experiments are shown, lines represent the means ± SDs. n_LB1/AurA_Prophase_ = 15 cells, n_LB1/PLK1_Prophase_ = 20 cells, n_AurA_Prophase_ = 17 cells, n_PLK1_Prophase_ = 17 cells, n_LB1_Prophase_ = 17 cells, n_LB1/AurA_Pro-meta/metaphase_ = 55 cells, n_LB1/PLK1_ Pro-meta/metaphase_ = 45 cells, n_AurA_ Pro-meta/metaphase_ = 47 cells, n_PLK1_ Pro-meta/metaphase_ = 54 cells, n_LB1_ Pro-meta/metaphase_ = 40 cells. *P* values were determined by one-way ANOVA with Tukey´s post hoc test.

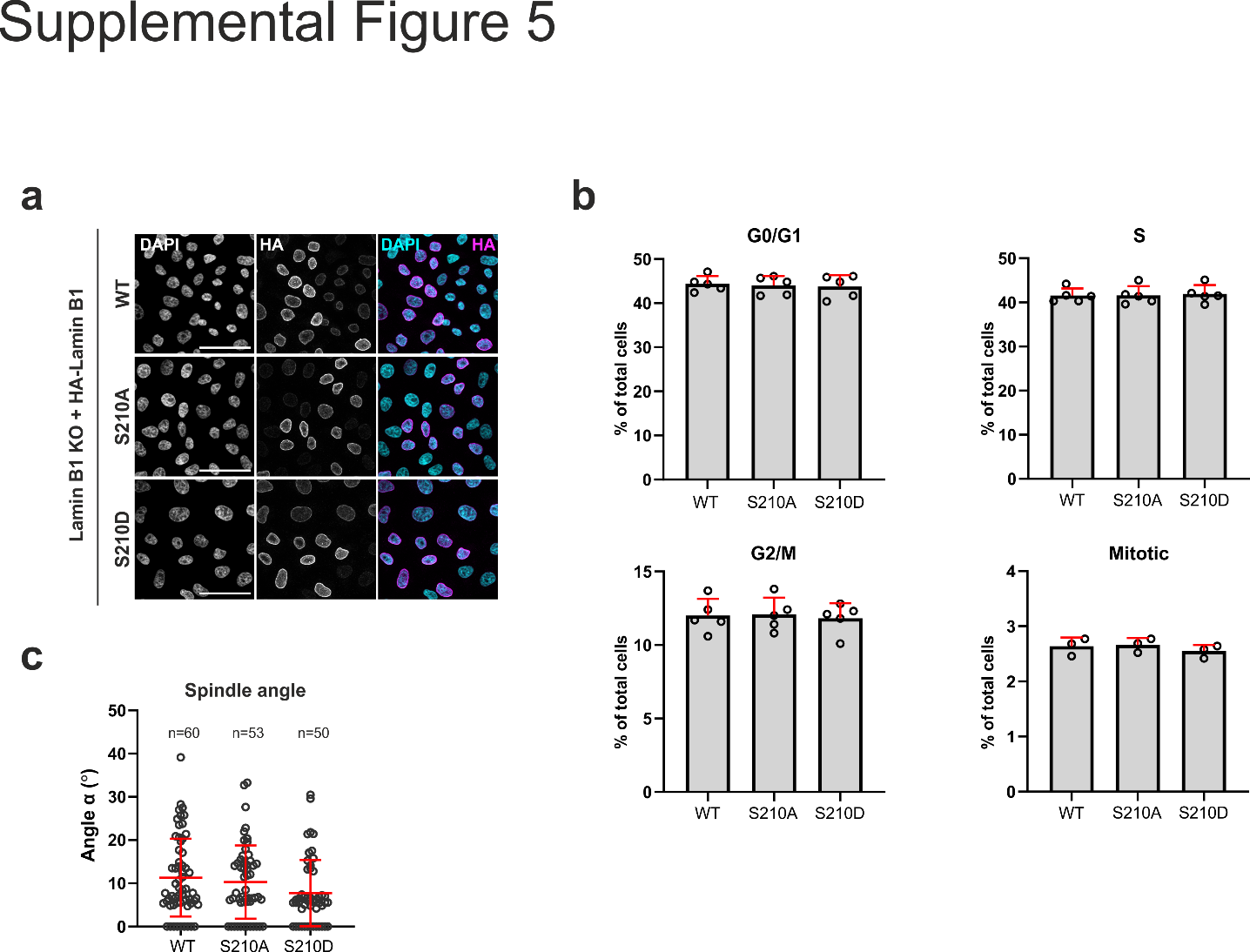

**Fig. S5: HA-Lamin B1 WT, S210A and S210D are expressed at similar levels and phosphorylation mutants do not affect cell cycle progression or spindle angle. a)** HeLa Lamin B1 KO cells reconstituted with HA-Lamin B1 WT, S210A or S210D were fixed and processed for immunofluorescence with anti-HA antibody. Nuclei were stained with DAPI. Scale bars, 10 μm. **b)** Reconstituted cells were processed for BrdU uptake and cell cycle analysis (panels G0/G1, S and G2/M) or for mitotic index determination (panel Mitotic) by flow cytometry. Data are mean + SD of 3-5 independent experiments. **c)** Quantification of spindle angle in each cell line. The angle was calculated as depicted in the scheme in Fig. 6c. The data presented are pooled from three independent experiments and the lines correspond to the mean ± SD. The total numbers of cells analyzed are shown in the graph.

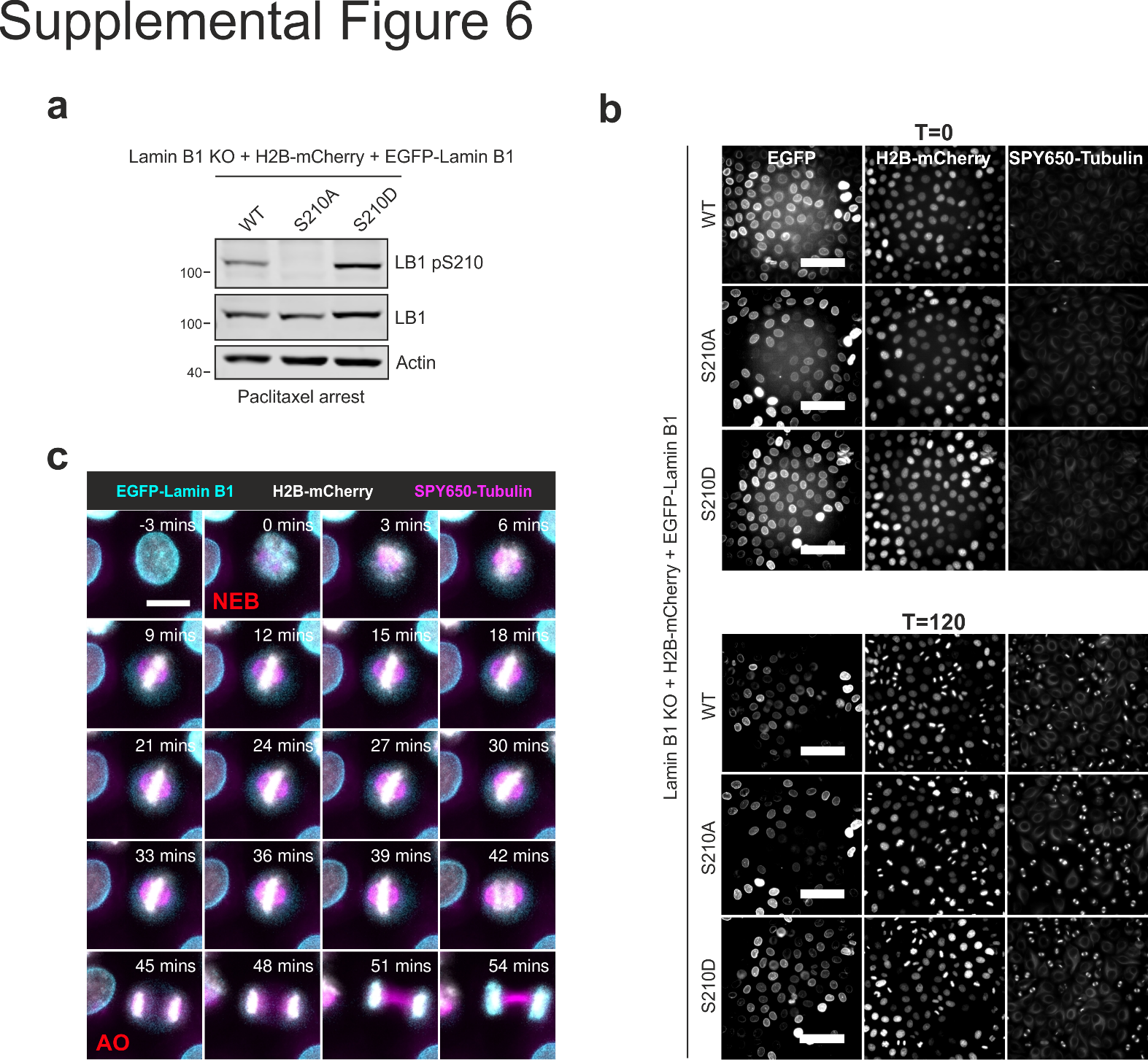

**Fig. S6: Live-cell imaging of Lamin B1 KO cells expressing H2B-mCherry and reconstituted with GFP-Lamin B1 WT, S210A or S210D. a)** Lysates from all three cell lines arrested with 100 nM paclitaxel for 18 hours were immunoblotted and probed with the indicated antibodies. **b)** Representative images of cell lines synchronized by double-thymidine block and stained with SPY650-Tubulin to visualize microtubules at time-point 0 (directly after the beginning of imaging, approx. 8 hours after release from the second thymidine block) and 120 minutes after. Scale bars, 100 μm. **c)** Example of the measurement of mitotic duration by live-imaging of a cell expressing EGFP-Lamin B1 WT. Mitotic duration was measured from the time of nuclear envelope breakdown (NEB) to anaphase onset (AO). Scale bar, 15 μm.

**Table S1:** ULK-dependent Lamin B1 phospho-sites detected by mass spectrometry.

| **Proteins** | **Position** | **Protein names** | **Loc. prob.** | **Score** | **Intensity LB1** | **Intensity ULK1** | **Intensity ULK3** | **Ratio ULK1/LB1** | **Ratio ULK3/LB1** |
| --- | --- | --- | --- | --- | --- | --- | --- | --- | --- |
| P20700 | 5 | Lamin-B1 | 1.00 | 138.08 | nd | nd | 0.347 | nd | >100 |
| P20700 | 19 | Lamin-B1 | 0.50 | 68.83 | 0.189 | nd | nd | <100 | <100 |
| P20700 | 20 | Lamin-B1 | 0.99 | 135.91 | 1.383 | 3.299 | 1.026 | 2.39 | 0.74 |
| P20700 | 23 | Lamin-B1 | 1.00 | 253.83 | 551.572 | 772.806 | 270.458 | 1.40 | 0.49 |
| P20700 | 28 | Lamin-B1 | 0.50 | 72.898 | nd | 0.035 | 0.866 | >100 | >100 |
| P20700 | 52 | Lamin-B1 | 0.50 | 64.878 | nd | nd | nd | nd | nd |
| P20700 | 55 | Lamin-B1 | 0.50 | 64.878 | nd | nd | nd | nd | nd |
| P20700 | 58 | Lamin-B1 | 0.93 | 99.566 | nd | 0.151 | 0.077 | >100 | >100 |
| **P20700** | **210** | **Lamin-B1** | **1.00** | **181.26** | **nd** | **0.232** | **0.103** | **>100** | **>100** |
| P20700 | 267 | Lamin-B1 | 1.00 | 129.16 | nd | 0.052 | 0.171 | >100 | >100 |
| P20700 | 278 | Lamin-B1 | 0.96 | 144.27 | 0.233 | 0.604 | nd | 2.59 | 0.00 |
| P20700 | 279 | Lamin-B1 | 0.96 | 110.39 | 0.233 | nd | nd | <100 | <100 |
| P20700 | 283 | Lamin-B1 | 0.60 | 101.53 | nd | 4.817 | nd | >100 | nd |
| P20700 | 284 | Lamin-B1 | 0.84 | 263.59 | nd | nd | 0.596 | nd | >100 |
| P20700 | 285 | Lamin-B1 | 0.84 | 85.457 | nd | 0.029 | 0.357 | >100 | >100 |
| P20700 | 288 | Lamin-B1 | 1.00 | 127.36 | nd | 0.025 | 0.029 | >100 | >100 |
| P20700 | 302 | Lamin-B1 | 1.00 | 82.543 | 0.071 | 0.270 | 0.071 | 3.80 | 0.99 |
| P20700 | 304 | Lamin-B1 | 0.49 | 77.593 | nd | nd | nd | nd | nd |
| P20700 | 305 | Lamin-B1 | 0.49 | 77.593 | nd | nd | nd | nd | nd |
| P20700 | 375 | Lamin-B1 | 1.00 | 66.262 | nd | 0.056 | nd | >100 | nd |
| P20700 | 520 | Lamin-B1 | 1.00 | 105.17 | nd | nd | 0.100 | nd | >100 |
| P20700;Q03252 | 391;405 | Lamin-B1;Lamin-B2 | 1.00 | 144.1 | 182.420 | 307.020 | 95.593 | 1.68 | 0.52 |
| P20700;Q03252 | 393;407 | Lamin-B1;Lamin-B2 | 1.00 | 123.28 | 113.799 | 237.174 | 68.452 | 2.08 | 0.60 |

GST-ULK1(1-649) or HIS-ULK3 were incubated with full-length Lamin B1 in the presence of kinase buffer and ATP. A reaction containing only Lamin B1 but no kinase (LB1) was used as control for phosphorylated sites present in the input substrate. The intensities of the phospho-peptides were normalized by the intensities of each protein. The ratio of the normalized values between the samples incubated with ULK1 or ULK3 and the substrate (Lamin B1) alone were calculated. Sites that were only detected in a specific sample were assumed to be at least 100-fold more abundant compared to the respective control sample. Sites with a localization probability (loc. prob.) higher than 0.75 were considered to be clearly localized (class I sites). Two ULK1-only (S283 and S375, depicted in blue), three ULK3-only (T5, S284, and S520, depicted in green) and six ULK1 and ULK3 (S28, S58, S210, S267, T285 and S288, depicted in red)-dependent phospho-sites were detected. The S210 site is highlighted in bold font. nd: not detected.

**Table S2**: Primers and vectors for recombinant and mammalian expression used in this study. Highlighted in grey are the constructs employed for experiments.

| **Nr #** | **Plasmid name** | **Primers** | **Cloning/**  **mutagenesis** | **Source** |
| --- | --- | --- | --- | --- |
| 1 | pET-20b(+)_HIS-LaminB1opt |  |  | Genscript |
| 2 | pGEX-Lamin B1_full | FOW 5´-TAAGGGATCCCCATGGCGACCGCGACCCCG-3´  REV 5´-AATGCGGCCGCTTCACATAATCGCGCAGCTACGGTTGCTC-3´ | Cloning of full Lamin B1 from plasmid # 1 into pGEX-5X-3  (NotI/BamHI) | This study |
| 3 | pGEX-Lamin B1 head (1-34) | FOW 5´-TGAAGCGGCCGCATCGTGAC-3´  REV 5´-CTCTTTCTCCTGCAGACGGCTCAGAC-3´ | Mutagenesis from plasmid # 2 | This study |
| 4 | pGEX-Lamin B1 coil 1 (35-215) | FOW 5´-TAAGGGATCCCCGAGCTGCGTGAACTGAAC-3´  REV 5´-AATGCGGCCGCTTCACTCTTCCTCGTACATGCTTTTAC-3´ | Cloning of Lamin B1 coil 1 from plasmid # 1 into pGEX-5X-3 (NotI/BamHI) | This study |
| 5 | pGEX-Lamin B1 coil 2 (216-386) | FOW 5´-TAAGGGATCCCCATTAACGAAACCCGTCGCAAG-3´  REV 5´-AATGCGGCCGCTTCATTCTTCTTCACCTTCCAGCAG-3´ | Cloning of Lamin B1 coil 2 from plasmid # 1 into pGEX-5X-3 (NotI/BamHI) | This study |
| 6 | pGEX-Lamin B1 Tail (387-586) | FOW 5´-CGTCTGAAGCTGAGCCCGAG-3´  REV 5´-GGGGATCCCACGACCTTCGATC-3´ | Mutagenesis of plasmid # 2 | This study |
| 7 | pGEX-Lamin B1 coil 1 (35-215)  S58A | FOW 5´-gcgGCGCTGCAGCTGCAAGTGAC-3´  REV 5´-GTTTTCGGTTTCCAGGCTACGAAC-3´ | Mutagenesis of plasmid # 4 | This study |
| 8 | pGEX-Lamin B1 coil 1 (35-215)  S210A | FOW 5´- gcgATGTACGAGGAAGAGTGAAGC -3´  REV 5´-TTTACGGAACTCCAGGTCTTC-3 | Mutagenesis of plasmid # 4 | This study |
| 9 | pGEX-Lamin B1 coil 2 (216-386) S267A | FOW 5´-gcgTACCATGCGAAGCTGGAGAAC-3´  REV 5´-TTGCTCCAGCTCCTCTTTATACAG-3´ | Mutagenesis of plasmid # 5 | This study |
| 10 | pGEX-Lamin B1 coil 2 (216-386) S284A | FOW 5´-gcgACCGTGAACAGCGCGCGTG-3´  REV 5´-GGTGTTCATTTCGCTGCTCAGACGCG-3´ | Mutagenesis of plasmid # 5 | This study |
| 11 | pGEX-Lamin B1 coil 2 (216-386) T285A | FOW 5´-gcgGTGAACAGCGCGCGTGAGG-3´  REV 5´-GCTGGTGTTCATTTCGCTGCTCAGAC-3´ | Mutagenesis of plasmid # 5 | This study |
| 12 | pGEX-Lamin B1 coil 2 (216-386) S288A | FOW 5´-gcgGCGCGTGAGGAACTGATGG-3´  REV 5´-GTTCACGGTGCTGGTGTTCATTTC-3´ | Mutagenesis of plasmid # 5 | This study |
| 13 | pGEX-Lamin B1 coil 2 (216-386) S284A/T285A/S288A | FOW 5´-AACgcgGCGCGTGAGGAACTGATGG-3´  REV 5´- CACcgccgcGGTGTTCATTTCGCTGCTCAGAC-3´ | Mutagenesis of plasmid # 5 | This study |
| 14 | pGEX-Lamin B1 coil 2 (216-386) S375A | FOW 5´-gcgGCGTACCGCAAACTGCTGG-3´  REV 5´-AATCTCCATATCCAGCGCCAG-3´ | Mutagenesis of plasmid # 5 | This study |
| 15 | pCMV6-Lamin B2 |  |  | Origene # RC200807 |
| 16 | pGEX-Lamin B2 coil 1 (49-229) | Insert PCR:  FOW 5´-TCGGATCTGATCGAAGGTCGTGGGATCCCCGAGCTTCGCGA  ACTGAACGATC-3´  REV 5´-AGATCGTCAGTCAGTCACGATGCGGCCGCTTCACTCCTCCTCG  AACACACTCTTC-3´  Vector PCR:  FOW 5´-AGCGGCCGCATCGTGACTGA-3´  REV 5´-GGGGATCCCACGACCTTCGATCAGATC-3´ | Cloning of Lamin B2 coil 1 from plasmid # 15 into pGEX-5X-3  (Gibson Assembly) | This study |
| 17 | pGEX-Lamin B2 coil 1 (49-229)  S224A | FOW 5´-GCAGTGTTCGAGGAGGAGTGAAG-3´  REV 5´-CTTCCGGAAGTCCAGCTCC-3´ | Mutagenesis of plasmid # 16 | This study |
| 18 | LaminB1-Myc/DDK |  |  | TWIST Biosciences |
| 19 | PCMV6_moULK1 (untagged) |  |  | Origene # MC206168 |
| 20 | Dendra2-Lamin B1-10 |  |  | Addgene_57728 |
| 21 | pMSCVpuro-mRFP-EGFP-rLC3 |  |  | *(77)* |
| 22 | pMSCVpuro-mRFP-EGFP-Lamin B1 WT | Insert PCR:  FOW 5´-GTACAAGTCCGGACTCAGATCTAGAGCGACTGCGACCCCCGT  GCCGCCGC-3  REV 5´- CTTTAGTTGGAAGTGGCTGTATGTCTGCTACATAATTGCACA  GCTTCTATTG-3´  Vector PCR:  FOW 5´-CAGACATACAGCCACTTCCAACTAAAG-3´  REV 5´-TCTAGATCTGAGTCCGGACTTGTAC-3´ | Cloning of Lamin B1 cDNA from plasmid # 20 into plasmid #21 to replace rLC3  (SLIC cloning). | This study |
| 23 | pMSCVpuro-mRFP-EGFP-Lamin B1 S210A | FOW 5´- gcaATGTATGAAGAGGAGATTAACGAGACC -3´  REV 5´- TTTGCGAAACTCCAAGTCCTC -3´ | Mutagenesis of plasmid # 22 | This study |
| 24 | pMSCVpuro-mRFP-EGFP-Lamin B1 S210D | FOW 5´- gacATGTATGAAGAGGAGATTAACGAGACC -3´  REV 5´- TTTGCGAAACTCCAAGTCCTC -3´ | Mutagenesis of plasmid # 22 | This study |
| 25-27 | pMSCVhygro-HA-Lamin B1 WT/S210A/S210D | FOW 5´-ATGTACCCATACGATGTGCCAGATTACGCCTCCGGACTCAGA  TCTAGAGCGAC- 3´  REV 5´-GGTGGCCGCTAGAACCTCGA-3´ | Mutagenesis of plasmid # 22-24 to replace mRFP-EGFP by HA, sub-cloning into pMSCVhygro | This study |
| 28-30 | pMSCVblast-mRFP-EGFP-Lamin B1 WT/S210A/S210D | Insert PCR:  FOW 5´-GCGCCGGAATTAGATCTCTCGAGGTTAACGATGGCCTTCTCCGAGGAC -3´  REV 5´- GGAAAAGCGCCTCCCCTACCCGGTAGAATTCTACATAATTGCACAGCTTCTATTGGATGC -3´  Vector PCR:  FOW 5´- AATTCTACCGGGTAGGGG -3´  REV 5´- CGTTAACCTCGAGAGATCTA -3´ | Subcloning of mRFP-EGFP-Lamin B1 WT/S210A/S210D from plasmid #22-24 into pMSCVblast (SLIC cloning). | This study |
| 31-33 | pMSCVblast-EGFP-Lamin B1 WT/S210A/S210D | FOW 5´- ATGGTGAGCAAGGGCGAGGAGCTG -3´  REV 5´- TAGATCTCTCGAGGTTAACG -3´ | Mutagenesis of plasmid # 28- 30 to remove mRFP | This study |
| 34 | pMSCVhygro_H2B-mCherry | H2B PCR:  FOW 5´- TCTCGAGCGTACGGCCGCCACCATGCCAGAGCCAGCGAAGTC -3´  REV 5´- ATGGTGGCGACCGGTGGATCCTTAGCGCTGGTGTACTTG -3´  mCherry PCR:  FOW 5´- CTAAGGATCCACCGGTCGCCACCATGGTGAGCAAGGGCGAGGAG -3´  REV 5´- AGAATTCGGGCCCTTACTTGTACAGCTCGTCCATGCCG -3´  Vector PCR:  FOW 5´- CTGTACAAGTAAGGGCCCGAATTCTACCGGGTAGGGGAGGC -3´  REV 5´- GACTTCGCTGGCTCTGGCATGGTGGCGGCCGTACGCTCGAGAG -3´ | Cloning of mCherry and H2B type 1-J amplified from cDNA into pMSCVhygro (Gibson assembly). | This study |

**Table S3:** Cell Profiler pipeline used for quantification of PLA signals in Fig.1b. Complete images were analyzed with the following pipeline.

| **Module** | **Detailed pipeline** |
| --- | --- |
| **Input** |  |
| **Images** | - Filter Images? 🡪 Images only  - Apply filters to the file list |
| **Metadata** | - Extract Metadata? 🡪 Yes  - Metadata extraction method 🡪 Extract from file/folder names  - Metadata source 🡪 File name  - Regular expression to extract from file name 🡪(?P<Cell>.*)_(?P<PLA>[A-C])_(?P<Picture>[0-9]^1^)_(?P<ChannelNumber>[0-3])  - Extract metadata from 🡪 All images  - Metadata data type 🡪 Text |
| **Names and Types** | - Assign a name to🡪 Images matching rules  - Process as 3D? 🡪 No  - Select the rule criteria 🡪 Match all of the following rules 🡪 Metadata Does HaveChannelNumber matching 🡪1  - Name to assign these images 🡪 rawDAPI  - Select the image type 🡪 Grayscale image  - Set intensity range from 🡪 Image metadata  - Select the rule criteria 🡪 Match all of the following rules 🡪 Metadata Does HaveChannelNumber matching 🡪2  - Name to assign these images 🡪 rawPLA  - Select the image type 🡪 Grayscale image  - Set intensity range from 🡪 Image metadata |
| **Processing** |  |
| **Identify Primary Objects: Nuclei** | - Use advanced settings? 🡪Yes  - Select input image 🡪 rawDAPI  - Name the primary objects to be identified 🡪 Nuclei  - Typical diameter of objects, in pixel units (Min,Max):100,300  - Discard objects outside the diameter range? 🡪Yes  - Discard objects touching the border of the image? 🡪Yes  - Threshold strategy 🡪Global  - Threshold method 🡪Otsu  - Two-class or three-class thresholding? 🡪Three classes  - Assign pixels in the middle intensity class to the foreground or the background? 🡪Background  - Threshold smoothing scale 🡪5  - Threshold correction factor 🡪1.2  - Lower and upper bounds on threshold 🡪0.02, 0.3  - Log transform before thresholding? 🡪Yes  - Method to distinguish clumped objects 🡪Shape  - Method to draw dividing lines between clumped objects 🡪Shape  - Automatically calculate size of smoothing filter for declumping? 🡪Yes  - Automatically calculate minimum allowed distance between local maxima? 🡪Yes  - Speed up by using lower-resolution image to find local maxima? 🡪Yes  - Display accepted local maxima? 🡪No  - Fill holes in identified objects? 🡪After both thresholding and declumping  - Handling of objects if excessive number of objects identified 🡪Continue |
| **Dilate Objects** | - Select the input object 🡪Nuclei  - Name the output object 🡪Nuclei_dilated  - Structuring element 🡪disk,10 |
| **Mask Image** | - Select the input image 🡪rawPLA  - Name the output image 🡪PLA_masked  - Use objects or an image as a mask? 🡪Objects  - Select object for mask 🡪Nuclei_dilated  - Invert the mask? 🡪No |
| **Identify Primary Objects: PLA foci** | - Use advanced settings? 🡪Yes  - Select input image 🡪 PLA_masked  - Name the primary objects to be identified 🡪 PLA_foci  - Typical diameter of objects, in pixel units (Min,Max):5, 40  - Discard objects outside the diameter range? 🡪Yes  - Discard objects touching the border of the image? 🡪Yes  - Threshold strategy 🡪Global  - Threshold method 🡪Otsu  - Two-class or three-class thresholding? 🡪Three classes  - Assign pixels in the middle intensity class to the foreground or the background? 🡪Background  - Threshold smoothing scale 🡪1.3  - Threshold correction factor 🡪0.5  - Lower and upper bounds on threshold 🡪0.04, 0.8  - Log transform before thresholding? 🡪Yes  - Method to distinguish clumped objects 🡪Intensity  - Method to draw dividing lines between clumped objects 🡪Intensity  - Automatically calculate size of smoothing filter for declumping? 🡪Yes  - Automatically calculate minimum allowed distance between local maxima? 🡪Yes  - Speed up by using lower-resolution image to find local maxima? 🡪Yes  - Display accepted local maxima? 🡪No  - Fill holes in identified objects? 🡪After both thresholding and declumping  - Handling of objects if excessive number of objects identified 🡪Continue |
| **Relate Objects** | - Parent objects 🡪Nuclei_dilated  - Child objects 🡪PLA_foci  - Calculate per-parent means for all child measurements? 🡪No  - Calculate child-parent distances? 🡪None  - Do you want to save the children with parents as a new object set? 🡪No |
| **Export To Spreadsheet** | - Select the column delimiter 🡪Tab  - Add a prefix to file names? 🡪No  - Overwrite existing files without warning? 🡪Yes  - Add image metadata columns to your object data file? 🡪No  - Add image file and folder names to your object data file? 🡪No  - Representation of Nan/Inf 🡪NaN  - Select the measurements to export 🡪Yes  - Data to export 🡪 Experiment 🡪 CellProfiler  🡪 Nuclei_dilated 🡪 Children 🡪 PLA 🡪 foci 🡪 Count  - Calculate the per-image mean values for object measurements? 🡪No  - Calculate the per-image median values for object measurements? 🡪No  - Calculate the per-image standard deviation values for object measurements? 🡪No  - Create a GenePattern GCT file? 🡪No  - Export all measurement types? 🡪Yes |

**Table S4:** Cell Profiler pipeline used for quantification of PLA signals in individual mitotic cells. Cells were manually scored and isolated, then analyzed with the following pipeline.

| **Module** | **Detailed pipeline** |
| --- | --- |
| **Input** |  |
| **Images** | - Filter Images? 🡪 Images only  - Apply filters to the file list |
| **Metadata** | - Extract Metadata? 🡪 Yes  - Metadata extraction method 🡪 Extract from file/folder names  - Metadata source 🡪 File name  - Regular expression to extract from file name 🡪^(?P<Cell>.*)_(?P<Cellcyclestage>[A-P])_(?P<Picture>[0-9]^1^)_(?P<ChannelNumber>[0-9])  - Extract metadata from 🡪 All images  - Metadata data type 🡪 Text |
| **Names and Types** | - Assign a name to🡪 Images matching rules  - Process as 3D? 🡪 No  - Select the rule criteria 🡪 Match all of the following rules 🡪 Metadata Does HaveChannelNumber matching 🡪2  - Name to assign these images 🡪 rawPLA  - Select the image type 🡪 Grayscale image  - Set intensity range from 🡪 Image metadata |
| **Processing** |  |
| **Identify Primary Objects: PLA foci** | - Use advanced settings? 🡪Yes  - Select input image 🡪 rawPLA  - Name the primary objects to be identified 🡪 PLA_dots  - Typical diameter of objects, in pixel units (Min,Max):5, 35  - Discard objects outside the diameter range? 🡪Yes  - Discard objects touching the border of the image? 🡪Yes  - Threshold strategy 🡪Global  - Threshold method 🡪Robust Background  - Lower outlier fraction 🡪 0.05  - Upper outlier fraction 🡪 0.05  - Averaging method 🡪 Mean  - Variance method 🡪 Standard deviation  - # of deviations 🡪 2.0  - Threshold smoothing scale 🡪1.1  - Threshold correction factor 🡪1.0  - Lower and upper bounds on threshold 🡪0.05, 1.0  - Method to distinguish clumped objects 🡪Intensity  - Method to draw dividing lines between clumped objects 🡪Intensity  - Automatically calculate size of smoothing filter for declumping? 🡪Yes  - Automatically calculate minimum allowed distance between local maxima? 🡪Yes  - Speed up by using lower-resolution image to find local maxima? 🡪Yes  - Display accepted local maxima? 🡪No  - Fill holes in identified objects? 🡪After declumping only  - Handling of objects if excessive number of objects identified 🡪Continue |
| **Export To Spreadsheet** | - Select the column delimiter 🡪Tab  - Add a prefix to file names? 🡪No  - Overwrite existing files without warning? 🡪Yes  - Add image metadata columns to your object data file? 🡪No  - Add image file and folder names to your object data file? 🡪No  - Representation of Nan/Inf 🡪NaN  - Select the measurements to export 🡪Yes  - Select measurments 🡪 Experiment 🡪 Metadata 🡪 Grouping Tags  🡪 Image 🡪 Count 🡪 PLA 🡪 dots  🡪 FileName 🡪 rawPLA  - Calculate the per-image mean values for object measurements? 🡪No  - Calculate the per-image median values for object measurements? 🡪No  - Calculate the per-image standard deviation values for object measurements? 🡪No  - Create a GenePattern GCT file? 🡪No  - Export all measurement types? 🡪No  - Data to export 🡪 Image  - Use the object name for the file name? 🡪 Yes |

**Table S5:** Image analysis pipeline (Harmony 5.2) for quantification of mitotic duration in EGFP-Lamin B1-expressing cells.

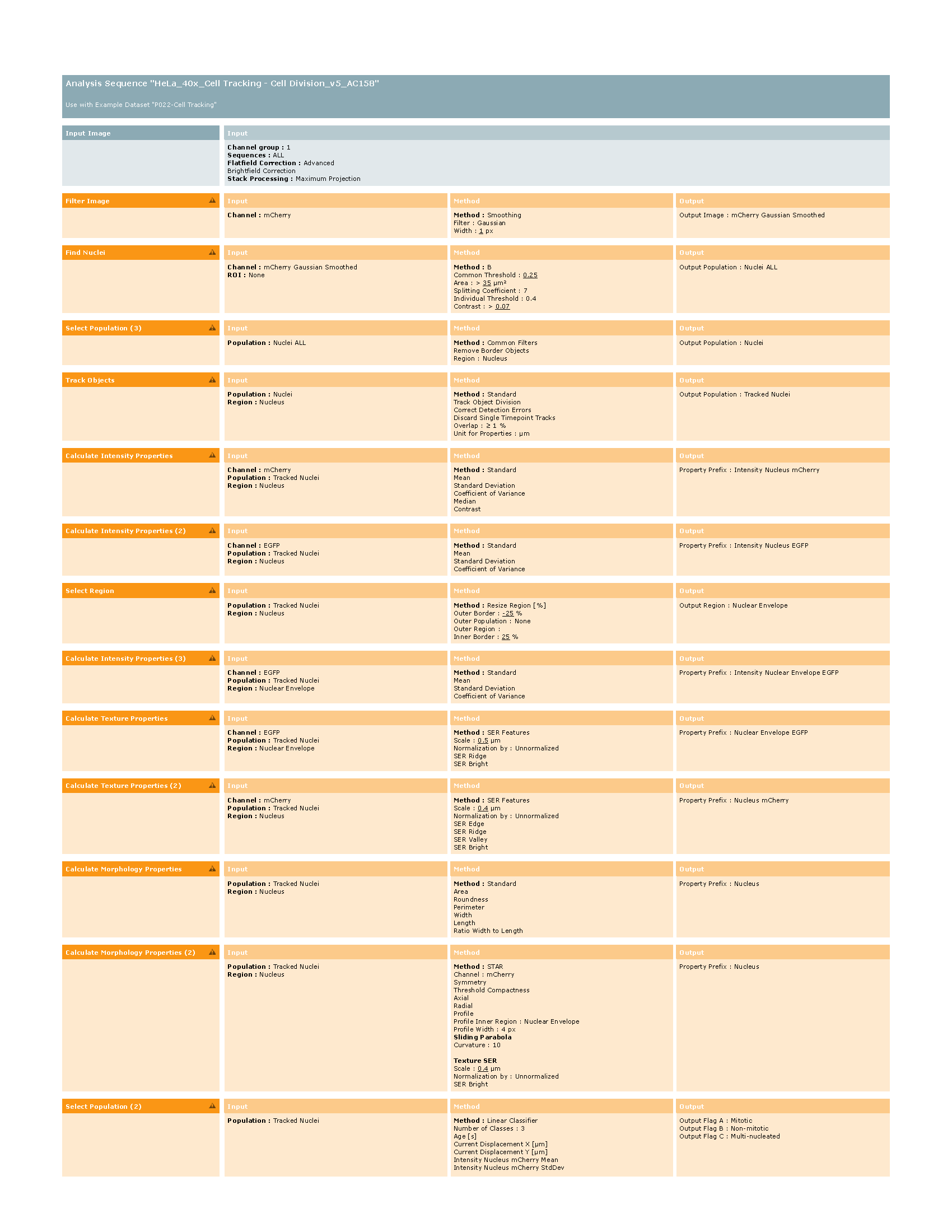

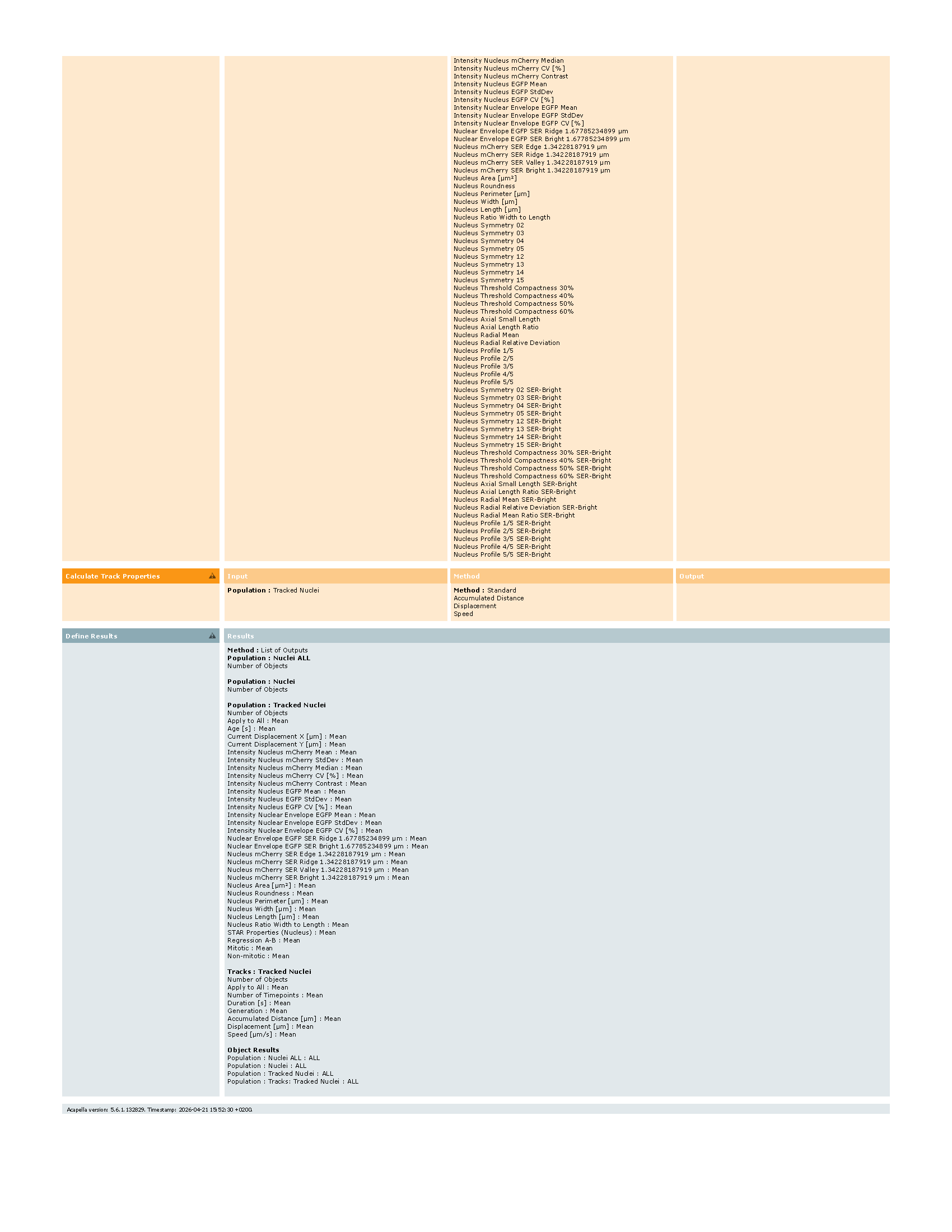

Data S1 (separate file): Lamin B1 p210 interactome. Lamin B1 p210 IPs were performed using lysates from paclitaxel (Pax)-arrested cells +/- λ phosphatase (each n=3). Shown are log2 transformed, median-normalized protein intensities. Proteins were shortlisted on being present in minimally 2 out of 3 replicates of Pax-arrested cells. Intensities of proteins being not detected in phosphatase-treated samples (Pax_L) were imputed using a normal distribution (width 0.3, down shift 1.3). Significant binders were determined using a one-sided Student`s t test (FDR<0.05) and are marked by a "+" in column G.
